# Copper oxide nanoparticles function as antineoplastic agents in uterine cancer cell lines

**DOI:** 10.64898/2026.06.07.729888

**Authors:** Jordan D. Berezowitz, Claire E. Rowlands, Lauren E. Mehanna, Breanna G. Knicely, Eva M. Goellner, Brittany E. Givens

## Abstract

Cancer of the uterine corpus is the fourth leading cancer and the fifth leading cause of cancer-related death in women in the United States. Chemotherapeutic resistance, specifically platinum-resistance, contributes to this problem. Therefore, an alternative treatment regimen is required. Using inorganic copper oxide nanoparticles (CuO NPs), we evaluated cancer cell responses indicative of anti-neoplastic activity. CuO NPs were characterized using transmission electron microscopy (TEM), scanning electron microscopy (SEM), energy dispersive spectroscopy (EDS), dynamic light scattering (DLS) and laser Doppler velocimetry (LDV). The nanoparticles were rod-like and had a diameter of 70 ± 30 nm and a copper content ranging from 77% - 82.6%. The hydrodynamic diameter and the zeta potential significantly decreased with more particles in solution. These materials were also used in four endometrial cancer cell lines and one cervical cancer cell line to evaluate cell viability, apoptosis, migration, and reactive oxygen species. In endometrial cancer cell lines, the IC50 values ranged from 1.028 ug/mL in HEC-1A cells to 73.62 ug/mL in Ishikawa cells, indicating that different cells have vastly different responses to CuO NPs. The results also indicated cell line-dependent differences in apoptosis, oxidation potential, and migration. Further, the cervical cancer cell line was modified using CRISPR technology to highlight a common germline mutation that causes earlier onset and more aggressive cancer progression. These genetic mutations resulted in differences in a loss of redox potential without observable changes in apoptosis or migration. The results of these studies indicate that CuO NPs elicit effects dependent upon the stage of cancer. Anticipated long-term applications of these studies includes the potential as a target-specific anti-cancer agent, designed using knowledge at the interface of colloids and the tumor environment.

**Graphical Abstract:** 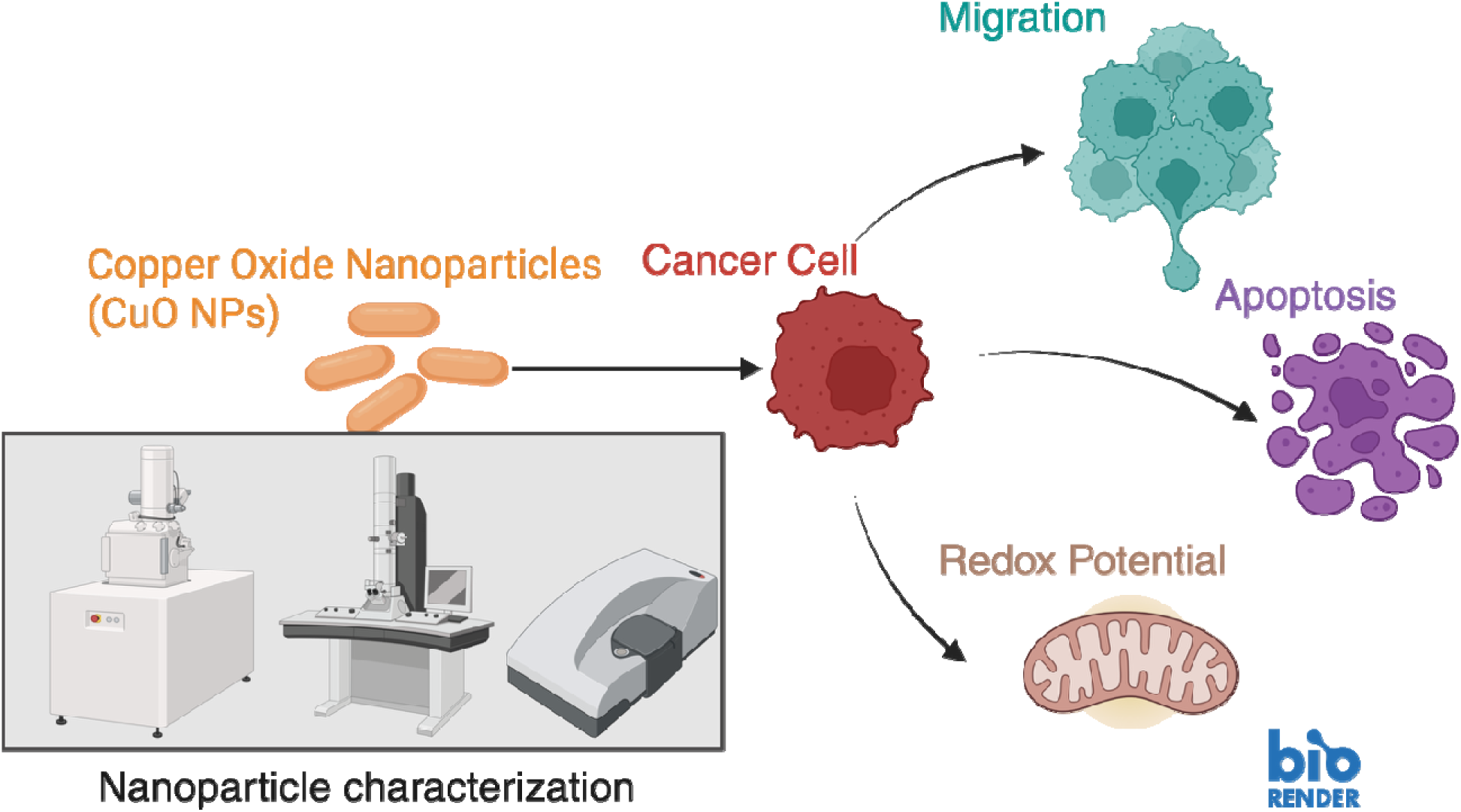

Copper oxide nanoparticles (CuO NPs) were characterized upon receipt using electron microscopy, elemental analysis, dynamic light scattering, laser Doppler velocimetry, and Fourier-transform infrared spectroscopy. These CuO NPs were exposed to HeLa cells with and without DNA mismatch repair deficiencies to assess the impacts on cancer cell migration, apoptosis, and redox potential.

## Introduction

The rising incidence of gynecological malignancies, including ovarian, endometrial, cervical, vaginal, and vulvar cancers presents a critical clinical challenge.^1^ Specifically, cancer of the uterine corpus (endometrial cancer) and uterine cervix (cervical cancer) have seen significant rises in incidence rates from annual increases of 0.6-1% and 1-2%, respectively.^1^ With over 34,000 gynecological cancer-related deaths projected in 2025 in the United States,^2^ there is a critical and urgent need for improved treatment strategies. The lack of early detection and screening mechanisms leaves surgery as the primary treatment option for endometrial cancer.^3–5^ Unfortunately, in many cases of advanced stage disease, surgery is not fully accessible or effective, and chemotherapy is also required.^3^ Despite that endometrial cancer is the most common gynecological malignancy worldwide, there are only two FDA-approved chemotherapeutic agents for endometrial cancer: Lenvatinib Mesylate (Lenvima) and Megestrol Acetate.^6^ As a result, some of the most powerful chemotherapeutics that are approved in ovarian cancer are used off-label in endometrial cancer, including cisplatin or carboplatin in combination with paclitaxel or doxorubicin.^7^ Platinum resistance in cancer chemotherapy is a major source of chemotherapeutic failure, leading to high recurrence and mortality.^8, 9^ Platinum resistance also reduces the efficacy of cisplatin or carboplatin; therefore, alternative strategies for effective chemotherapy are required.

Platinum resistance arises from enhanced efflux, neutralization of reactive oxygen species, or efficient DNA-damage repair. In uterine cancer, heritable DNA mismatch repair (MMR) mutations from Lynch Syndrome additionally result in earlier age at diagnosis (oncogenesis) and more aggressive tumors (disease progression). Lynch syndrome is a set of heritable germline mutations in the *MLH1, MSH2, MSH6,* or *PMS2* genes. These genes impair DNA-damage repair pathways, which leads to cancer progression. Lynch syndrome occurs in 30 – 40% of individuals diagnosed with endometrial cancer,^10, 11^ and therefore is a critical consideration in developing potential endometrial cancer treatments.

To bypass these mechanisms of resistance, inorganic metal oxide nanoparticles (MeOx NPs) offer a promising therapeutic alternative. MeOx NPs have unique, size-dependent physicochemical properties, a high surface-area to volume ratio, and elevated density of active surface sites compared to the bulk counterparts with significantly different biological reactivity.^12, 13^ The nanotoxicity of MeOx NPs has been widely investigated across various biological interfaces, including the pulmonary system,^14, 15^ in model organisms such as zebrafish,^16^ and the intestines.^17–20^ Within MeOx NPs, copper (II) oxide nanoparticles (CuO NPs) are known to be toxic to the lungs and intestines, believed to be mediated by reactive oxygen species (ROS), a primary cause of oxidative stress.^21^ In tumor cells, ROS can induce apoptosis and disrupt tumor growth.^22^ Therefore, we hypothesized that CuO NPs could be used as a therapeutic material in tumor cells.

In this study, the potential of CuO NPs as an alternative cancer therapeutic for endometrial cancer was assessed. Four endometrial cancer cell lines were used, and to further investigate the effects on genetic mutations common in endometrial cancer, CRISPR-modified HeLa cells were also incorporated (**Table 1**). The breadth of these cell lines enabled investigations into the antineoplastic potential of CuO NPs in endometrial cancer, with respect to stage and grade. Additionally, using the CRISPR-modified cells, the studies probed DNA mutations common in Lynch syndrome.^23^

**Table 1.**
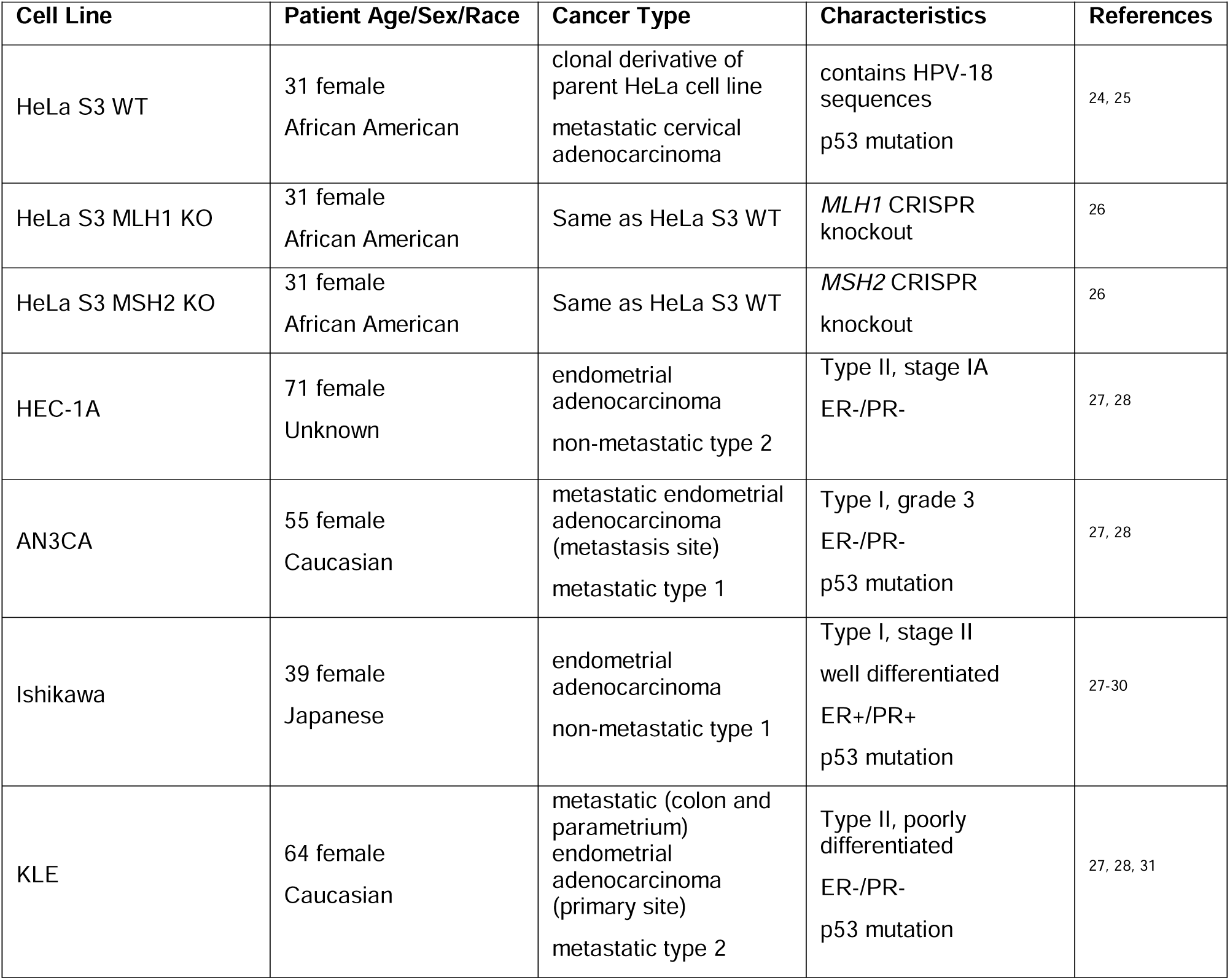
Cell lines used in this study and the stage, grade, tissue, and gene mutations associated with each. Abbreviations: clustered regularly interspaced short palindromic repeats (CRISPR), estrogen receptor negative (ER-), estrogen receptor positive (ER+), progesterone receptor negative (PR-), progesterone receptor positive (PR+).

## Materials and Methods

### Materials

Copper (II) oxide nanoparticles, copper (II) nitrate trihydrate (Cu(NO_3_)_2_·3H_2_O), Dulbecco′s Phosphate Buffered Saline (DPBS), penicillin-streptomycin (PS), dimethyl sulfoxide (DMSO), were purchased from Sigma-Aldrich (St. Louis, MO). Dulbecco’s Modified Eagle Medium (DMEM), Dulbecco’s Modified Eagle Medium/Nutrient Mixture F-12 (DMEM:F12), Thiazolyl Blue tetrazolium bromide (MTT), ethanol, Alexa fluor 647 phalloidin, eBioscience3 Annexin V apoptosis detection kits FITC and propidium iodide, Prolong Diamond antifade mountant with DAPI, rat tail collagen I, and Trypsin-EDTA (0.25%) were purchased from Thermo Fisher Scientific (Waltham, MA). Fetal bovine serum (FBS) was purchased from Bio-Techne (Minneapolis, MN). Isopropanol (IPA) was purchased from VWR (Radnor, PA). Lacey carbon 300 mesh nickel-coated TEM grid was purchased from Ted Pella (Redding, CA). GSH/GSSG-Glo™ Assay kit was purchased from Promega (Madison, WI).

### Copper oxide characterization

Scanning electron microscopy (SEM) (FEI Helios NanoLab 660, FEI Company, Hillsboro, OR) was used to analyze particle size and morphology. A 1 mg/mL solution of CuO NPs was prepared in ultrapure DI water and deposited with either no sonication or 60 minutes of sonication. Sonication was performed in a Fisher Scientific FS30 water bath sonicator at 130W ultrasonic power and 40kHz frequency. A 10 µL aliquot of the CuO NP solution was added as a single drop to silicon wafers atop SEM stubs. The water evaporated under ambient conditions, leaving a dry sample. The samples were coated with approximately 2.5 nm of platinum using a Lecia Ace 600 carbon/sputter coater (Lecia Microsystems, Durham, NC). SEM was performed at 2.0 kV accelerating voltage, 25.0 pA current, and 15,000X magnification. Particle size was quantified for 250 particles across multiple fields of view using ImageJ ^32^.

Transmission electron microscopy (TEM) (FEI Talos F200X, FEI Company, Hillsboro, OR) was conducted to obtain complementary electron micrographs. Microscopy was performed after sonicating a 1 mg/mL solution of CuO NPs in ethanol for either 10 minutes or 60 minutes. Following sonication, 10 µL of the solution was pipetted onto a 300 mesh nickel-coated lacey carbon TEM grid. The ethanol was then evaporated under ambient conditions for 10 minutes before imaging. In conjunction with TEM, energy-dispersive X-ray spectroscopy (EDS) was performed on the CuO NPs to determine the copper and oxygen content. Elemental composition from nickel-coated lacey carbon grids were also quantified.

Colloidal characterization was conducted through laser Doppler velocimetry (LDV) and dynamic light scattering (DLS) of CuO NPs in both PBS and ultrapure DI water at room temperature with and without sonication. DLS was used to measure the hydrodynamic diameter (D_H_) of CuO NPs. To collect triplicate measurements without sonication, stock solutions of 1 mg/mL CuO NPs suspended in either 1X DPBS and ultrapure DI water were used to create 1 mL dilutions of desired concentrations of 10 and 20 µg/mL. After vortexing for 10 seconds, solutions were transferred to a clear cuvette and placed in the Malvern Zetasizer Nano ProBlue (Malvern Panalytical, Westborough, MA). Three independent samples of each concentration and dispersant type were measured three times each with a 60 second equilibrium time between sample measurements. For sonicated samples, the 1 mg/mL stock solution was placed in a Fisher Scientific FS30 water bath sonicator at 130W ultrasonic power and 40kHz frequency for 60 minutes before performing dilutions to a desired concentration. To measure zeta potential a disposable capillary zeta cell was filled with 900 µL of either 10 or 20 µg/mL of CuO NP in 1X DPBS. The capillary zeta cell was capped on both ends and placed in the Malvern Zetasizer Nano ProBlue at room temperature (25°C). Triplicate measurements on each of the three independent solutions were measured. The capillary zeta cell was flushed with ultrapure DI water between each sample.

### Cell maintenance

This study used four endometrial carcinoma cell lines: KLE, Ishikawa, AN3CA, and HEC-1A, and one cervical cancer cell line, HeLa S3, with two CRISPR-modified knockouts (KOs): HeLa S3 *MLH1* knockout, and HeLa S3 *MSH2* knockout. Each cell line was cultured under specific conditions and passaged at a confluency optimized for the cell lines. KLE, AN3CA, and HEC-1A cells were purchased from American Type Culture Collection (ATCC, Manassas, VA) and cultured in DMEM-F12 supplemented with 10% (v/v) fetal bovine serum (FBS) and 1% (v/v) penicillin-streptomycin (PS). Ishikawa H cells were gifted from Dr. Aliasger Salem (University of Iowa, Iowa City, IA) and cultured in DMEM supplemented with 10% (v/v) FBS and 1% (v/v) PS. HeLa S3 and HeLa S3 knockouts were cultured in RPMI-1640 containing 10% (v/v) FBS and 1% (v/v) PS. All cell lines were cultured in T75 flasks in a 37°C humidified incubator at 5% CO_2_ and passaged at 90% confluency, with media refreshed every two days.

HeLa *MLH1* KO and HeLa *MSH2* KO cell lines were generated with single guide RNA (sgRNA) CRISPR-Cas9 for *MLH1* and *MSH2* as previously published in Miller *et al*. ^26^. Briefly, the LentiCRISPRv2 was a gift from Feng Zhang (Addgene plasmid #52961), and the plasmid was digested with BsmBI and gel purified using the QIAquick PCR purification kit according to the manufacturer’s instructions. Complementary oligonucleotides (Integrated DNA Technologies, Coralville, IA) encoding the sgRNA were then annealed and cloned into LentiCRISPRv2. Cells were then transfected with Lipofectamine 3000 (Thermo Scientific L3000008), and the cells were selected with puromycin (Promega, Madison, WI). Single cell clones were allowed to grow up under puromycin selection and expanded. Loss of MLH1 and MSH2 protein expression was confirmed for each clone using SDS-PAGE and western blot analysis.

### Cell viability and IC50 concentration

Relative cell viability of treated cells compared with an untreated control were assessed using MTT assays, and the results from this study were used to calculate an IC50 value for each cell line. Cells were seeded in 96-well plates at 4.69 x 10^5^ cells/cm^2^. The next day, cells were treated with 12 half-log concentrations of 60-minute sonicated CuO NPs ranging from 0 to 1000 µg/mL of CuO NPs or 0 to 3000 µg/mL of copper nitrate trihydrate for 24 hours.

MTT solution was prepared by mixing a 5 mg/mL solution of Thiazolyl Blue tetrazolium bromide, 98% powder in sterile 1X DPBS, protected from light. The stopping solution was prepared using 90% isopropyl alcohol with 10% (v/v) DMSO. MTT solution was adding to cell media at a 1:10 ratio. Cells were washed twice with 100 µL of 1X DPBS prior to being treated with 110 µL of MTT solution in cell media and incubated for 4 hours. All media containing MTT solution was removed, then 100 µL of stopping solution was added to each well to solubilize the formazan dye. Plates were shaken lightly on a Vortex Genie and centrifuged at 1000 rcf (Eppendorf 5810R, Hamburg, Germany) for 10 minutes to remove CuO NPs from suspension. The supernatant was transferred to a new clear 96-well plate before being read using a Synergy H1 microplate reader (Agilent Technologies, Santa Clara, CA) at an absorbance of 570 nm.

### Oxidative Stress

Clear bottom, white-walled, tissue-culture treated 96-well plates were coated with 10 µg/cm^2^ Rat Tail, Collagen I for one hour at ambient room temperature (∼24°C) to enhance cell attachment. Collagen solution was removed, and plates were washed once with 150 µL of 1X DPBS. Cells were plated at 4.69 x 10^5^ cells/cm^2^ for optimum luminescence measurement and sufficient cell counts. Cells were incubated at 37°C and 5% CO_2_ and left to attach overnight. Media was removed, and cells were treated with respective cell media containing 0, 10, or 20 µg/mL of CuO NPs. The treatments were performed in triplicate wells for both total glutathione and oxidized glutathione measurements. After 24 hours, the media was removed, and cells were washed once with 150 µL 1X DPBS. The GSH/GSSG-Glo™ Assay kit (Promega) was used following the manufacturer’s instructions. Background luminescence of the reagent was determined using cell-free wells, which was subtracted from the luminescence measurements of treated cell samples.

### Cell migration assay

An Ibidi 2-well silicone insert with a defined 500 µm cell-free gap between the wells was placed in each well of a 6-well culture plate. Each cell line was seeded at 1.36 x 10^5^ cells/cm^2^ in their respective cell media and incubated at 37°C and 5% CO_2_ overnight to allow for cell adherence. Once a confluent cell monolayer was present within each well of the insert, the insert was removed and the wells were washed with 1X DPBS to remove any floating cells, then replaced with the appropriate serum-free cell media. Serum-free media was supplemented containing 1% (v/v) penicillin-streptomycin and exchanged every 24 hours after the cell-free gap was created.

To test the effect of CuO NPs on cell migration, a 1 mg/mL CuO NP stock solution was prepared in 1X DPBS and sonicated for 60 minutes. The stock solution was then diluted to create a working solution of 10 µg/mL or 20 µg/mL CuO NPs in the appropriate serum-free cell media for each cell line. After the culture insert was removed from the plate, the well was washed with 1X DPBS and replaced with the 10 µg/mL or 20 µg/mL CuO NP working solution. The cells were exposed to the CuO NP working solution for 24 hours, then exchanged with serum-free media without CuO NPs every 24 hours for the remainder of the timepoints. Images of the cell-free gap were taken on an Eclipse Ti-E (Nikon, Melville, NY) inverted microscope at 0, 6, 24, and every 24 hours thereafter for 240 hours or until the gap was closed. ImageJ MRI Wound Healing Tool was used to determine the gap area at each time point for representative images from n = 6 independent samples for each cell type and treatment group ^33^.

### Cell apoptosis and necrosis

Each cell line was seeded at 2.37 x 10^5^ cells/cm^2^ in a 6-well plate and incubated at 37 °C and 5% CO_2_ overnight. Cell media was removed, and 3 mL of CuO NP solutions were added at 10 µg/mL and incubated for 24 hours. After 24 hours, CuO NPs and cell media were removed, and the cells were washed with 3 mL of 1X DPBS. Then, 1 mL of 0.25% Trypsin-EDTA was added to each well. Cells were incubated in Trypsin-EDTA at room temperature until about 90% of the cells detached from the plate. 2 mL of complete media was added to each well to neutralize trypsin, and cells were collected into individual 15 mL centrifuge tubes. Cells were centrifuged at 500 rcf for 3 minutes to obtain pellets. These were resuspended, washed with 3 mL of 1X DPBS, and centrifuged under the same conditions to form a cell pellet. The supernatant was discarded, and cells were suspended in 1 mL of 1X Binding Buffer diluted from the 10X stock using ultrapure DI water. Cell counts were obtained for all tubes and verified that they contained over 1 million cells.

All CuO NP-treated cell tubes and one of the control tubes were resuspended in 500 µL of binding buffer, and 25 µL Annexin V staining solution was added and mixed. These cells were incubated at ambient room temperature (∼25°C) for 15 minutes before being centrifuged to form a pellet. The supernatant was discarded, and cells were washed with 2 mL of Binding Buffer, centrifuged to form a pellet with supernatant discarded. The treated cell tubes were suspended in 500 µL of 1X Binding Buffer and the Annexin V control was suspended in 750 µL of Binding Buffer. CuO-treated cells received 12.5 µL of PI staining solution and were placed on ice. Untreated cells were used as a blank control, suspended in 1 mL of 1X Binding Buffer, and placed on ice. One vial of untreated cells was used as the Propidium Iodide (PI)-stained control and were suspended in 750 µL of 1X Binding Buffer, to induce cell death for a positive control, this centrifuge tube was placed in a water bath at 65 °C for 15 minutes, then 18.75 µL of PI solution was added to the dead cell solution. 500 µL of this solution was added to a new tube and placed on ice. 500 µL of the Annexin V control were added to a new tube and placed on ice. 250 µL of the Annexin V control was added to a new tube and combined with 250 µL of the PI-stained control. This Annexin V and PI-stained cell solution was used as the combined control. Prepared solutions were analyzed using a BD FACSymphony^TM^ A5 Cell Analyzer in the Flow Cytometry and Immune Monitoring Core Shared Resource Facility. All three replicate treatments and controls were analyzed at 50,000 counts for consistency.

### Intracellular copper (ICP-MS)

Each cell line was seeded at 5.26 x 10^4^ cells/cm^2^ in a 6-well plate and incubated at 37 °C and 5% CO_2_ overnight. Cell media was removed, and 3 mL of CuO NP solutions were added at either 10 µg/mL or 20 µg/mL and incubated for 24 hours. After 24 hours, CuO NPs and cell media were removed, and the cells were washed with 3 mL of 1X DPBS. Then, 750 µL of 0.25% Trypsin-EDTA was added to each well. Cells were incubated in Trypsin-EDTA at room temperature until about 90% of the cells detached from the plate. 1.5 mL of media was added to each well to neutralize trypsin, and cells of identical treatment wells were collected into metal-free 15 mL centrifuge tubes. Cell pellets were obtained with centrifugation, and the supernatant was discarded. Cells were then resuspended and washed twice with 3 mL of 1X DPBS and centrifuged to form a pellet. After the second wash, the supernatant was discarded, and the cells were frozen at -80 °C for 24 hours before lyophilization. Cell pellets were weighed before and after lyophilization to obtain wet and dry masses. Data are reported as µg Cu/g dry cell mass, and fold changes in copper content were calculated using the mean µg Cu/g dry cell mass for samples in triplicate.

## Results

Copper oxide nanoparticles (CuO NPs) were purchased from Sigma-Aldrich and used without further modification. The manufacturer’s specifications indicated that nanoparticles were less than 50 nm in diameter and 77.0-82.6% copper. These dimensions were assessed using SEM (**Figure 1A**), TEM (**Figure 1B**), and EDS (**Figure 1C**). The particles were rod-like; therefore, the longest dimension was taken as the particle diameter: 70 ± 30 nm, and the copper composition ranged from 81% to 87% across different samples, calculated from EDS elemental quantification (**Figure S1**).

**Figure 1.**
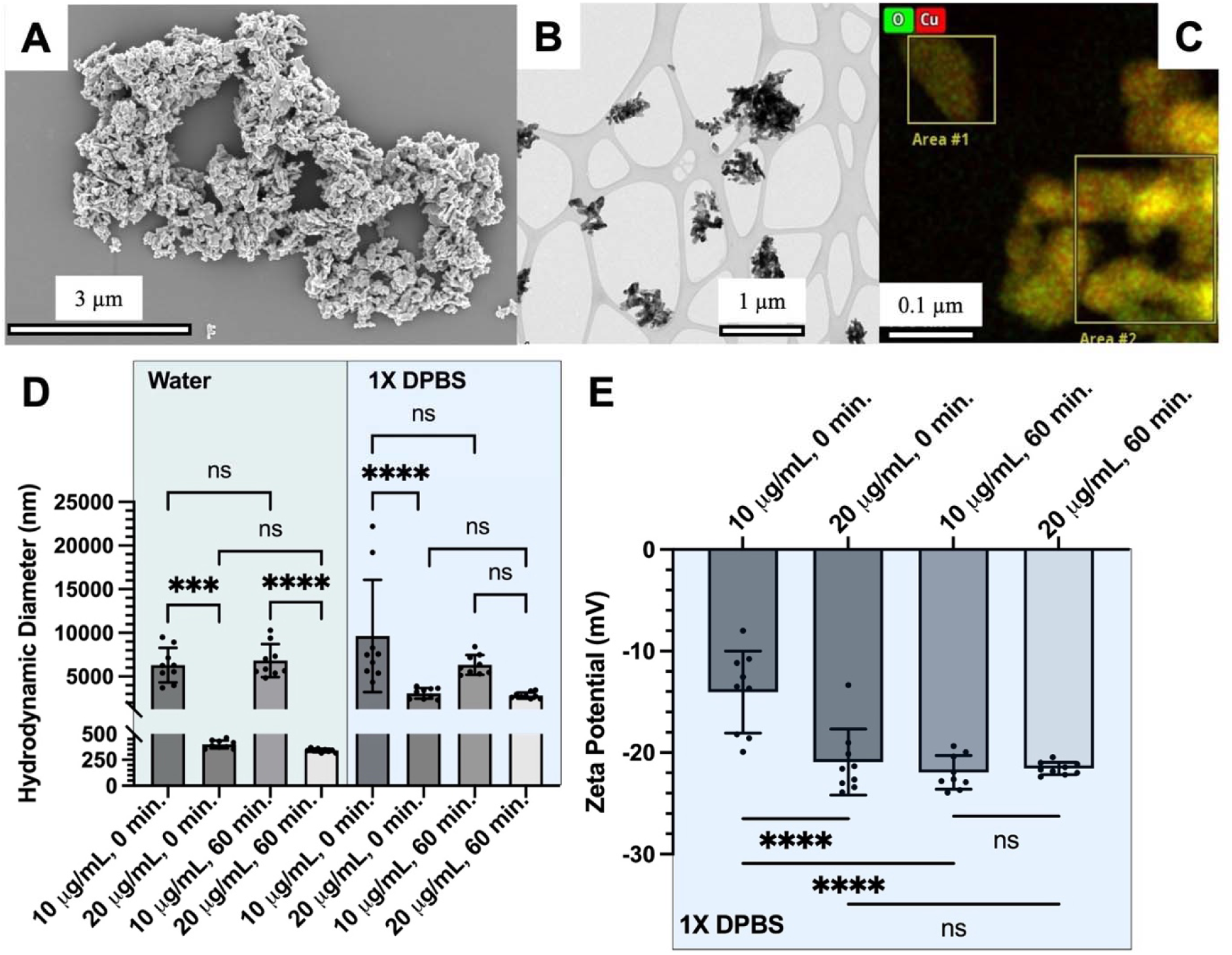
Copper oxide nanoparticle characterization with (a) SEM, (b) TEM, (c) EDS, (d) DLS, and (e) laser Doppler velocimetry.

The colloidal stability was assessed in deionized water and 1X DPBS (**Figure 1D**). 1X DPBS was used as a dispersant to simulate physiological conditions with respect to pH and salinity. The resultant hydrodynamic diameter indicated particle aggregation in both solutions, with greater aggregation in 1X DPBS than in ultrapure DI water. Significant decreases in aggregation were noted between the different concentrations of CuO NPs at each sonication time, but no significant differences were noted between the same concentration at different sonication times in DI water. In 1X DPBS, however, the only significant decrease in aggregation occurred at higher concentration without sonication. The zeta potential measurements were conducted only in 1X DPBS to meet conductivity requirements (**Figure 1E**). The zeta potential decreased significantly (p < 0.0001) for the 10 μg/mL concentration from -21.94 ± 1.668 to -14.04 ± 4.045 upon sonication. Additionally, in the absence of sonication, the zeta potential significantly decreased (p < 0.0001) with increasing concentration to -20.93 ± 3.256. In contrast, there were no significant differences (p = 0.9907) for the 20 μg/mL concentration upon sonication, nor were there significant differences (p = 0.9612) between 10 μg/mL and 20 μg/mL for 60 minutes sonication. These differences in zeta potential indicate greater the best colloidal stability for a 10 μg/mL suspension without any sonication.

Quantified cell uptake of copper was measured by ICP-MS using the total amount of intracellular copper per dry cell mass (**Figure 2**). There were no significant changes (p > 0.05) in uptake by dose in any of the cell lines. Even so, the Ishikawa cells internalized significantly more copper at 20 µg/mL than all other cell lines (p < 0.01 for KLE cells and p < 0.0001 for all other cell lines). Quantified particle uptake in each cell line was determined by inductively coupled plasma mass spectrometry (ICP-MS). ICP-MS results indicate that there is a dose-dependent relationship for uptake for all endometrial and healthy human cell lines. Cervical cancer cell lines (HeLa S3 WT, *MLH1*, and *MSH2* KOs) showed an increase in uptake from the control (0 µg/mL), but there was a lower copper content in the 20 µg/mL treatment than the 10 µg/mL treatment. All cell lines showed increased levels of intracellular copper after exposure to CuO NPs, indicating that copper is being internalized by cells at concentrations as low as 10 µg/mL. The fold change increase in copper content between the 10 and 20 µg/mL treatment groups for each respective cell line was analyzed (**Table S1**). The largest fold-change increase occurred in Ishikawa cells at 9.46 ± 3.72 and 3.48 ± 1.67, respectively. All three HeLa S3 cell lines showed a less than one-fold-change, indicating a decrease in intracellular copper content from 10 to 20 µg/mL treatments. All other cell lines showed between a one- to two-fold-change between treatments, which would be as expected based on a linear trend in uptake.

**Figure 2.**
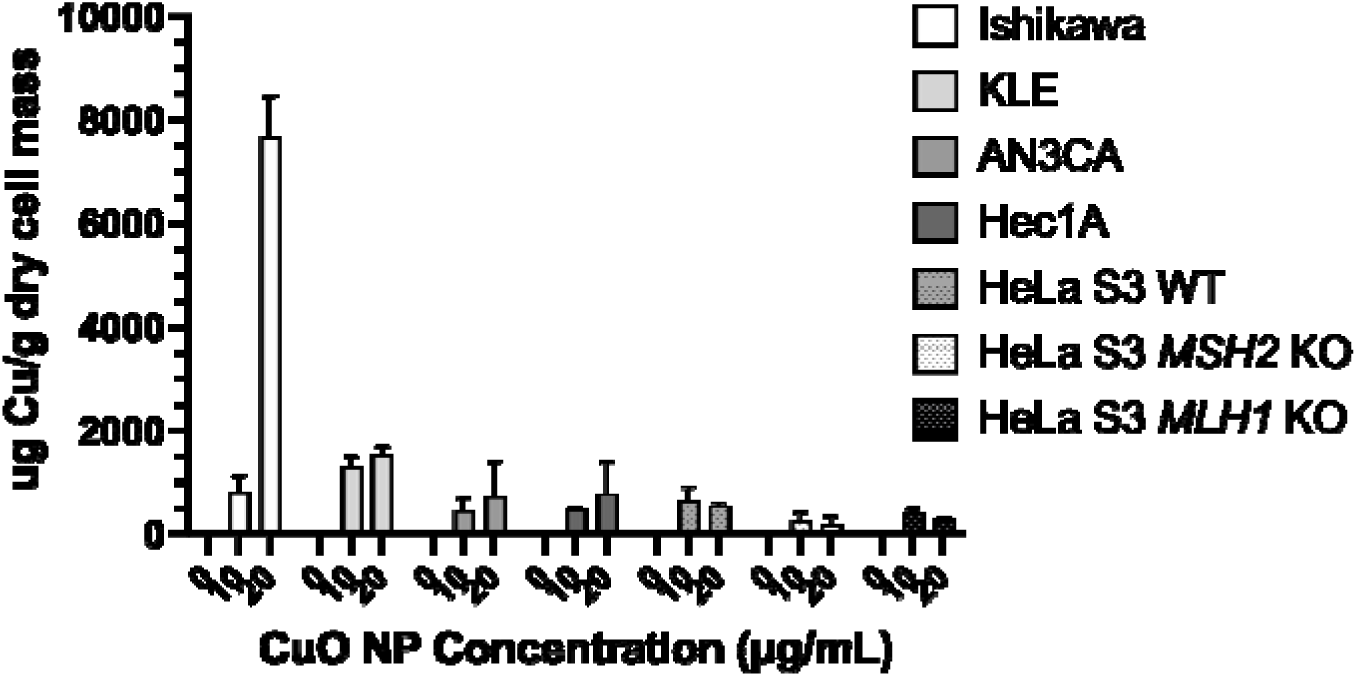
Quantified uptake of CuO NPs via ICP-MS following 24 hours of exposure *in vitro*.

The IC50 values for each cell line were determined using the MTT assay to assess mitochondrial activity (**Figure 3**). The HEC-1A cells had the lowest IC50 value of 1.028 µg/mL. Complete cell death was measured at concentrations of 10 µg/mL and greater with sensitivity occurring at 0.1 µg/mL. The largest dose-dependent response to CuO NPs occurred in AN3CA and Ishikawa cells, with sensitivity occurring at concentrations greater than 0.01 µg/mL, with complete cell death occurring at concentrations above 100 µg/mL, with the most gradual decrease in relative viability indicated by the slope of the dose-response curve. Larger IC_50_ values were calculated for the KLE cell line at 72.40 µg/mL. Sensitivity to CuO NPs occurred at concentrations above 10 µg/mL, and relative cell viability above 25% was observed at 100 µg/mL, which was not observed in any other cell line. Complete cell death was observed in concentrations above 100 µg/mL. KLE cells represent the least sensitive cell line when exposed to CuO NPs. The calculated IC50 values for the parent HeLa S3, HeLa S3 MLH1 KO, and HeLa S3 MSH2 KO cell lines were 11.23, 13.93, and 14.46 µg/mL, respectively. Complete cell death was observed for all HeLa S3 cell lines for concentrations at and above 100 µg/mL and sensitivity to CuO NPs occurred at concentrations at or above 1 µg/mL.

**Figure 3.**
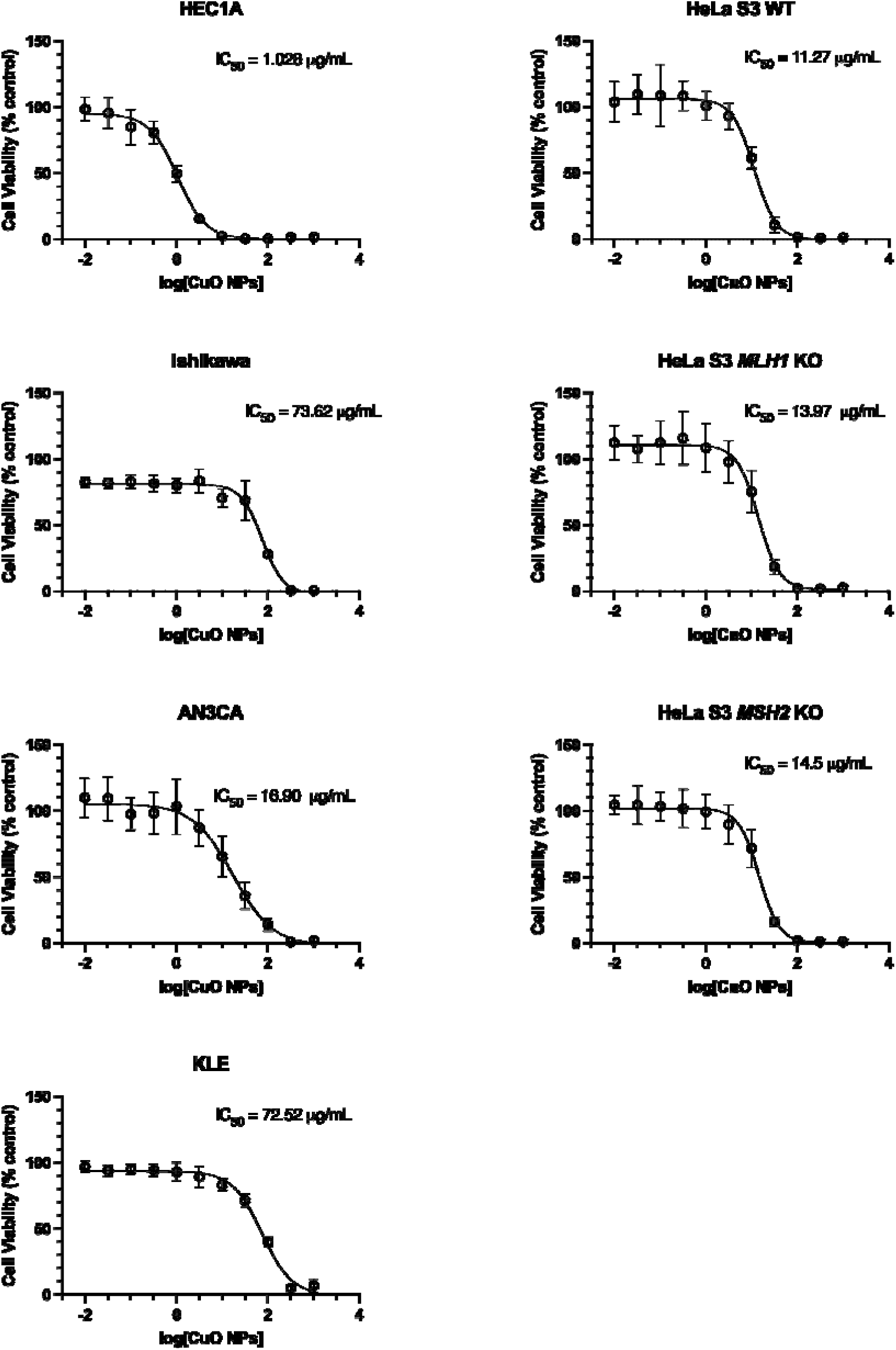
Minimum inhibitory concentration (IC50) following 24 hours of CuO NP exposure *in vitro*.

Cancer cell apoptosis, migration and reactive oxygen species were used to evaluate the antineoplastic potential of CuO NPs (**Figure 4**). Apoptosis and necrosis were determined using an Annexin V/propidium iodide stain and flow cytometry. The KLE cells had the greatest proportion of live cells following 10 μg/mL CuO NPs, with an average of 87.7% live cells, and less than 5% of cells in each of the other phases of cell death. The Ishikawa cells had 53.3% live cells with 29.0% of cells in early apoptosis, 11.0% of cells in late apoptosis and 6.7% of cells in necrosis. The AN3CA cells had 41.3% of live cell with 54.8% of cells in early apoptosis and only 1.8% and 2.1% of cells in late apoptosis and necrosis, respectively. The HEC1A cell line was the most sensitive with just 9.8% live cells and 82.3% of cells in early apoptosis. In this cell line, less than 1% of cells were in necrosis and the remaining 7.2% were in late apoptosis. Cancer cell apoptosis was also determined in the HeLa S3 WT, HeLa S3 *MLH1* KO, and HeLa S3 *MSH2* KO. These cells all had relatively high proportions of live cells, with the WT having 84.3%, the *MLH1* KO having 83.2% and the *MSH2* KO cells having 86.6% live cells. Of the remaining 15% of cell in each cell line, the *MLH1* KO cells had the greatest amount of cells in apoptosis with 11.4% in late apoptosis and 3.6% in early apoptosis. The WT cells had the greatest proportion in necrosis, with 5.0%, and approximately 5% (4.4% and 6.3%) of cells in early and late apoptosis, respectively. The *MSH2* KO cells were primarily in early apoptosis with 7.1% of cells in this phase; 2.9% of cells in late apoptosis and 3.4% of cells in necrosis

**Figure 4.**
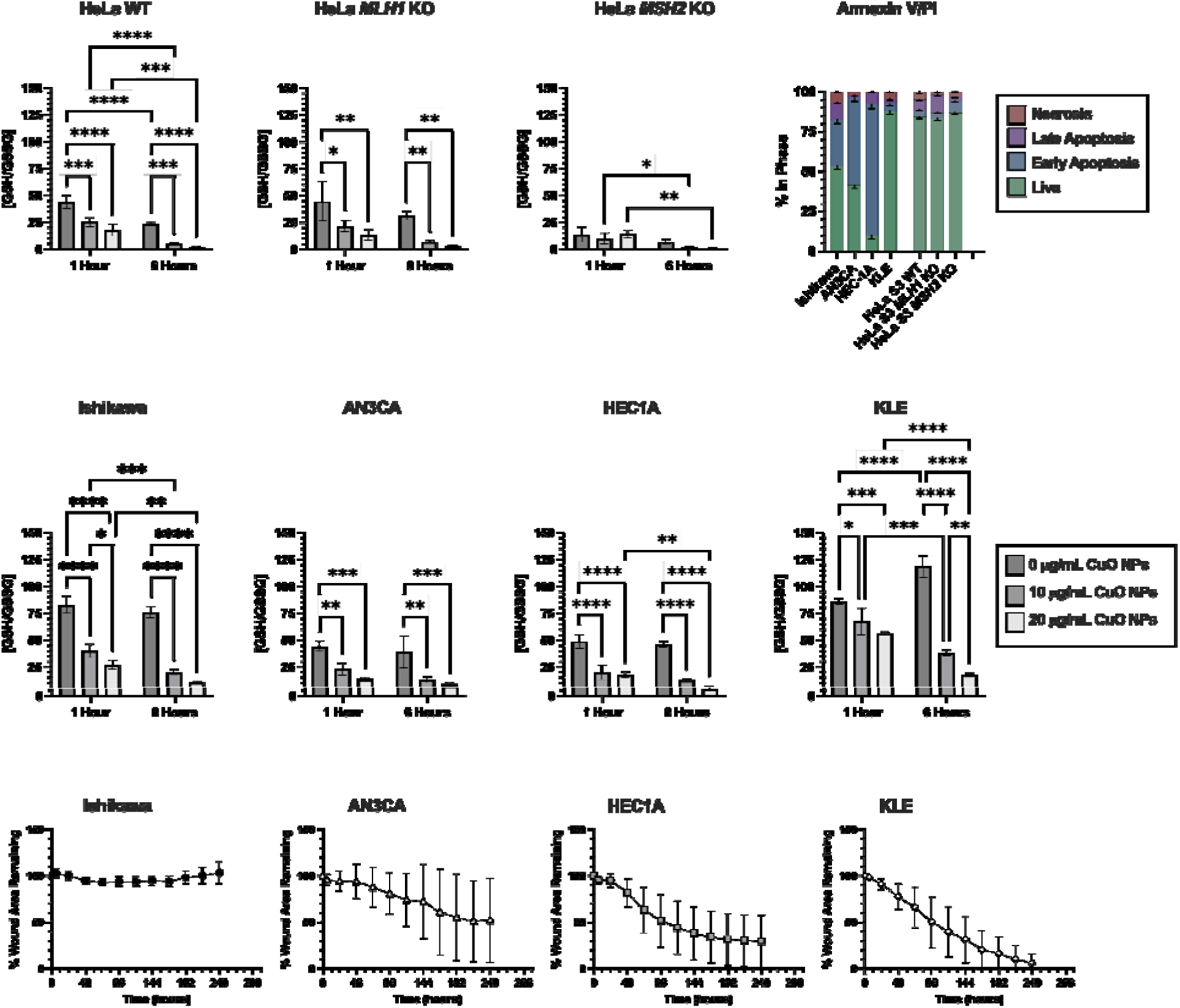
Cytotoxicity of CuO NPs as assessed via GSH/GSSG balance, Annexin V/PI staining and flow cytometry, and wound healing (scratch) closure. Data are provided for all cell lines, except in the case of the scratch assay in which the HeLa cells did not have quantifiable results.

The wound healing (scratch assay) for migration potential of the cancer cells with 10 µg/mL of CuO NPs indicated that the Ishikawa cells increased the wound area over time, whereas the AN3CA, HEC1A, and KLE cells all decreased the wound area (**Figure 4**). Would healing was assessed in an untreated control **(Figure S2**), 10 µg/mL of CuO NPs (**Figure S3**), and 20 µg/mL of CuO NPs (**Figure S4**). In HeLa cells, all copper concentrations induced cell detachment, preventing quantification of the wound-healing area. The KLE cells showed the greatest wound closure over seven days, followed by HEC1A and AN3CA. Over seven days, the Ishikawa cells did not close any of the wound area following exposure to 10 µg/mL CuO NPs. The 20 µg/mL CuO NP exposure was not quantified (**Fig. S4**) due to excessive cell death, hypothesized to result from impaired cell-cell signaling.

Excess ROS generation and oxidative stress were the hypothesized mechanisms of toxicity for CuO NPs. Therefore, the cellular response to oxidative stress was quantified through the natural antioxidant defense system of glutathione ^34^. The relative ratio of reduced (GSH) to oxidized forms of glutathione (GSSG) was quantified in each cell line (**Figure 4**). The baseline oxidative potential was determined using the highest GSH/GSSG ratio at 0 µg/mL at 1 and 6 hours. KLE cells had the highest baseline oxidative potential, and after one hour of exposure, KLE cells showed a sustained antioxidant response with the highest GSH/GSSG ratios recorded across all cell lines. Ishikawa cells had greater oxidative stress levels than HEC-1A the AN3CA, and all HeLa S3 cell lines. In the HeLa cells, the MSH2 KO cells had significant differences in the mean GSH/GSSG ratios when comparing treated (10 and 20 µg/mL) to the control (0 µg/mL) at matched time points. The difference in the HeLa MSH2 KO cell line is attributed to the lowest baseline antioxidant potential, compared to all other cell lines, which show significant oxidative stress at baseline and at 6 hours of exposure.

## Discussion

Chemotherapy for advanced-stage cancer is necessary to shrink tumors either before or after surgery, but often inefficient due to chemotherapeutic resistance ^8, 9^. Alternative strategies for inducing cell death in cancer cells may be achieved by combating the hallmarks of cancer as a guide ^35^. This project aimed to assess these hallmarks through cell death, proliferation, and invasion following exposure to a novel anti-neoplastic agent, copper oxide nanoparticles. These particles have been shown to exhibit toxic properties and, therefore, can be harnessed therapeutically. Furthermore, because copper is an essential micronutrient, residual copper at low concentrations may be less harmful to healthy tissues than platinum accumulation from platinum-based chemotherapeutics commonly used to treat this disease. An in vitro exploration of the effects of CuO NPs on gynecological cancers across a variety of cell lines will provide a strong screening of the potential of this material as a therapeutic.

The characterization of commercially available copper (II) oxide nanopowder was performed in both 1X DPBS and pure ultrapure DI water, mimicking the suspension within an aqueous biological environment. Upon the addition of either dispersant to create a 1 mg/mL solution, aggregates of particles were observed. After vortexing for 30 seconds and leaving the suspensions undisturbed, particles precipitated out of the solution and settled at the bottom of the vial. With goals of distributing particles throughout the solution and preventing aggregates from remaining in solution, a water bath sonicator was used to provide ultrasonic waves at a high frequency to agitate particles through cavitation bubbles ^36^. Cavitation causes mechanical agitation that results in enhanced mixing and the breakdown of aggregates into smaller sizes or individual particles ^37^. In SEM, particles were not sonicated as the surface scan of SEM was able to render quality micrographs. In contrast, with TEM CuO NPs were sonicated for 10 minutes to disperse aggregates as necessary for the transmission beam to render quality micrographs.

The aggregation was further quantified through the hydrodynamic diameter (D_H_) using DLS. Stock solutions of 1 mg/mL of CuO NPs in 1X DPBS and pure ultrapure DI water were sonicated for 60 minutes and used to create 10 and 20 µg/mL dilutions for DLS. In water, the hydrodynamic diameter significantly decreased with and without sonication as the particle concentration increased, indicating that the higher particle concentration increased the colloidal stability. However, this effect was not observed in 1X DPBS, which was likely a result of the presence of salts on colloidal stability.^38^ Optimizing sonication time resulted in adequately distributed particles and reduced agglomeration of particles within solution at 60 minutes. Hydrodynamic diameter measurements for dilutions of 10 and 20 µg/mL created in ultrapure DI water were significantly lower than identical concentrations in 1X DPBS. This phenomenon can be attributed to the thermodynamic properties of colloidal systems ^39^. Zeta potential measurements remained consistent for both 0- and 60-minute sonication times, implying the net charge of particles is unaltered upon sonication. However, the variation of individual measurements was reduced with increasing sonication time at matched concentrations. Therefore indicates that the zeta potential was more stable after sonication.

Aggregates of 60-minute sonicated particles in 1X DPBS were approximately 3 µm in hydrodynamic diameter and -22 mV for zeta potential. This is likely due to the reduced agglomeration effectively stabilizing surface charge after sonication. Based on this characterization and exploration into the effect of sonication on CuO-NP agglomerates and zeta potential, 60-minute sonication of 1 or 10 mg/mL CuO NPs in 1X DPBS was used prior to all dilutions made for cell culture experiments. These properties are not idea for cellular uptake due to the cell electrostatic repulsion due to the negative charge of the cell membrane, and the large size making internalization by the cell limited ^40^. The hypothesized mechanism for uptake of particles of this size is macropinocytosis or surface-associated binding ^41^. Results of ICP-MS analysis of lysed dry cell pellets revealed increases in copper content from the controls (0 µg/mL), indicating that copper was internalized or bound to the surface of cells. The results indicate different amounts of copper internalized for each cell line. Results indicated that Ishikawa cells had the greatest uptake capacity, as evidenced by the large amounts of copper observed in cells treated with 20 µg/mL. Prior research indicated that KLE and Ishikawa cells had similar trends in uptake of PLGA nanoparticles and hypothesized that the KLE cells have higher expression of efflux transporters than Ishikawa cells, explaining the differences in uptake magnitude ^42^. The expression of efflux transporters, as well as the mechanism of uptake, such as micropinocytosis, can be used to further explain these differences ^43, 44^. Metallic nanoparticles with strong reductive or oxidant power elicit cytotoxicity through ionic and/or electronic transfers ^45^. These transfers occur during dissolution, oxidation-reduction, and catalytic reactions in cell medium.

The HeLa S3 cells with knockouts of the DNA mismatch repair proteins MLH1 and MSH2 had greater IC50 values than the parent cell line, indicating a lower toxic response to CuO NPs.

In endometrial cancer cells, the IC50 of cisplatin ranges from 0.022 – 0.56 _µ_g/mL (730.7 – 1,859 _µ_M) and carboplatin from 0.096 – 1.20 _µ_g/mL (258.6 - 3232 _µ_M) ^46^. These values are well above the ideal pharmacological range for reducing off-target effects, as evidenced by the widespread side effects of chemotherapy. In contrast, the IC50 values for CuO NPs ranged from 12.95 – 925.8 _µ_M, indicating that this may be a better therapeutic in certain cell lines capable of reducing existing off-target effects.

Among the endometrial cancer cell lines, HEC-1A cells exhibited the lowest IC_50_ value (1.028 µg/mL), indicating that this early stage at isolation may increase CuO sensitivity. HEC1A cells also exhibited complete cell death at the lowest concentration compared with all other cell lines. AN3CA cells exhibited the second-lowest IC_50_ value (16.90 µg/mL), followed by approximately equal values for the KLE cells (72.52 µg/mL) and the Ishikawa cells (73.62 µg/mL). Although the low IC50 for the low stage and grade HEC1A cells is promising for early-stage cancers as an alternative to hysterectomy, which is the current first-line treatment, the absence of a trend among the other cancer cell lines according to stage, grade, type, and metastatic stage, indicates that further investigations are required to understand CuO NP sensitivity. For example, KLE cells heightened expression of efflux transporters, which may be responsible for the decreased response ^42^.

HeLa S3 WT and CRISPR-modified knockouts all showed similar trends to one another, indicating that the gene knockouts did not influence MTT-related cell viability. The results indicated that CuO NPs are more anti-neoplastic to early-stage, less aggressive forms of endometrial cancer than metastasized cervical and endometrial cancer cell lines, potentially pointing to differences in their mechanism of toxicity. In one study, A549 cells exposed to 50 µg/mL of CuO NPs for 24 hours showed a significant increase in MSH2 protein expression, involved in mismatch repair of double-stranded DNA breaks, compared to untreated cells ^47^. This points to the resistance to toxicity observed in HeLa S3 MSH2 KO cells compared to WT and MLH1 KO cells. Additionally, A549 cells had increased expression of p53 protein and Hsp70, which indicate a stress response to the CuO NP exposure ^47^.

Results showed that KLE cells had the highest percentage of live cells among all cell lines. This aligns with the viability data showing that at 10 µg/mL of CuO NPs cells are not experiencing significant toxicity. Conversely, the HEC-1A and the AN3CA cells showed the lowest number of cells in the live state, indicating that these cells were more sensitive to CuO NPs, further supporting the viability results. Both cell lines also showed a significant number of cells in early apoptosis, indicating that CuO NPs are inducing apoptosis as their mechanism of cell death. Further exploration into specific caspase proteins can determine what is activating this apoptosis pathway ^48^. Flow cytometry was used to assess apoptosis and necrosis. Annexin V is a calcium-dependent protein that binds to phosphatidylserine (PS), which is localized to the extracellular membrane only during apoptosis. PS is also activated by certain caspase cascades, such as caspase 3 and caspase 8, which can provide further mechanistic insights into CuO NP-mediated apoptosis ^49, 50^. In fact, caspase 3 is a confirmed pathway for CuO NP-mediated apoptosis in several cell lines ^51^, indicating that Annexin V is a robust metric for CuO NP apoptosis.

Mitochondrial respiration is the primary source of intracellular reactive oxygen species generation ^52, 53^. KLE cells exhibited the lowest oxidative stress, which may be related to the high observed IC50 value compared with other cell lines. These data indicate that oxidative stress may be the mechanism of toxicity to the mitochondria, ultimately showing that antioxidant potential may be preventing the cell from triggering the apoptosis pathway and KLE cells continue to proliferate. The IC50 is an important consideration in pharmacology and therefore when considering a potential alternative cancer treatment. An IC50 value in the nanomolar (nM) or micromolar (_µ_M) range is indicative of few off-target effects.

For example, the KLE cells, which are metastatic endometrial cancer cells and have a *BRCA2* mutation were the least sensitive to CuO NPs ^30^. This may indicate that *BRCA2* mutations do not enhance responses to CuO NPs despite these mutations enhancing platinum sensitivity ^54^. Additionally, the Ishikawa cells, which also had a high IC50 value, have mutations in both *BRCA1* and *BRCA2* ^30^, further suggesting that BRCA mutations do not enhance copper sensitivity. Additionally, the HeLa MLH1 and MSH2 knockouts exhibited higher IC50 values than the HeLa WT. These data indicate that despite the accepted mechanism of action for CuO NP-induced cell death, mutated DNA damage repair genes do not enhance CuO NP sensitivity in these gynecological cancer cell lines. Comparing these IC50 values with the intracellular copper concentration reveals that, despite high uptake in Ishikawa cells, the IC50 remains high. In comparison, the HeLa S3 KOs had the lowest intracellular copper, including less than the HeLa S3 WT at the same dose. Together, these data indicate that internalization efficiency, defined as the percentage of dosed CuO NPs internalized, is not predictive of cell death.

Further examination of the effects on cell death and other cancer cell characteristics, including migration and oxidative stress in the endometrial cancer cells and the cervical cancer cells. Previous research has shown that ROS are generated during incubation of CuO NPs in IEC-6 rat epithelial cells, thereby causing cytotoxicity ^19^. In primary brain astrocytes, concentrations of CuO NPs above 64 µg copper per mL (1000 µM), there was an increase in LDH activity, and decreased cell viability by MTT. At low concentrations of CuO NPs (100 µM and below) weak ROS staining of cells was observed after 3 hours; however, for concentrations at 300 or 1000 µM CuO NPs, strong ROS staining was observed after 3 hours of treatment ^55^. Additionally, Cu^2+^ ions leached from a Cu_2_O core in Cu_2_O/Au-Pt@MOF@F127 particles were effective at increasing ROS-mediated toxicity in the tumor microenvironment of glioblastoma, confirmed via GSH depletion ^56^. These particles contained 90.9% Cu^2+^ and 9.1% Cu^+^ ^57^. These studies further bring to light that many mechanisms of toxicity may be at play for CuO-NP interaction with cells.

In endometrial cancer cells, KLE cells had the highest baseline oxidative stress potential, followed by Ishikawa cells. This high baseline potential for reactive oxygen species may be related to the high IC50 values, as the leading mechanism of action for CuO NP-related cell death is the generation of reactive oxygen species. The ability of the KLE and Ishikawa cells to combat this reactive oxygen potential may make them more resistant to CuO treatment, as evidenced by high IC50 values and high intracellular localization of copper. In contrast, the HEC1A and AN3CA cells, which had lower I50 values, had lower baseline oxidative potential and had lower GSH/GSSG ratios following 6 hours of 20 μg/mL treatment compared with the Ishikawa and KLE cells. Furthermore, the proportion of live KLE and Ishikawa cells following 10 μg/mL CuO NP exposure indicates that the cells are inducing apoptosis following the CuO treatment as readily as the other cell lines. Although the AN3CA cell line had approximately 40% live cells, the highest percentage of the cells were in early apoptosis. The HEC1A cell line had over 80% of cells in early apoptosis and only 10% of cells alive. These data correspond with the IC50 values where HEC1A cells had the lowest IC50 value and the lowest proportion of living cells following treatment well above the IC50. Interestingly, the migration assay indicated that Ishikawa cells did not possess any migration potential after treatment with 10 ug/mL CuO NPs whereas the KLE cells had the greatest migration potential after treatment. These data indicate that cancer cell migration, an indicator of metastatic potential, may be governed by the stage at which cells were isolated rather than by treatment. The AN3CA and HEC-1A both had slowed migration potential compared with the control, and neither cell line closed the entire wound area over seven days. Although the HeLa S3 cells all had high survival according to the apoptosis assay, the shift from necrosis in the WT cells to apoptosis in the mismatch repair knockout cells may indicate that these pathways indeed play a role in cell replication and survival.

Endometrial, colon and ovarian cancers, are linked to Lynch syndrome; genetic mutations in either *MLH1* or *MSH2*, which impact DNA mismatch repair mechanisms. Individuals with Lynch syndrome often have earlier onset of endometrial cancer compared to those without these gene mutations ^10, 58, 59^. To evaluate the influence of Lynch syndrome on CuO NP therapy for cancer, we used CRISPR-Cas9 to knockout either MLH1 or MSH2 in HeLa S3 cells, resulting in deficient MLH1 and MSH2 proteins, which was confirmed using western blot ^26^. These results indicated that the MSH2 KO had a lower baseline GSH/GSSG ratio compared with the WT and MLH1 KO, which had similar GSH/GSSG baseline ratios. Additionally, as the concentration of CuO NPs increased, there were no significant decreases in the GSH/GSSG ratio. However, this may be due to the lower baseline, as the MSH2 KO had the lowest GSH/GSSG ratio across all cell lines after 6 hours (**Figure 4**). For all HeLa S3 cells, there was a large proportion of live cells (∼85%) remaining at the time of analysis. The MSH2 KO cells had the greatest proportion of cells in early apoptosis, and the MLH1 KO had the greatest proportion of cells in late apoptosis. These results indicate that indeed the DNA damage repair KO influenced the mechanism of cell death; however, further investigation is required to determine how this is mediated in cells. These HeLa S3 cells are semi-adherent; therefore, the migration assay could not be performed, as cell attachment was significantly impaired in this assay ^60^.

## Conclusion

This study provides a scoping analysis of the effects that CuO NPs have on endometrial and cervical cancer cells *in vitro*, providing further direction to specific mechanistic studies to investigate the varied antineoplastic effects CuO NPs have on gynecological tissues. This study is the first to investigate the genotoxic and anti-neoplastic properties of CuO NPs *in vitro* using multiple endometrial and cervical cancer cell lines across differentiation status, stage, and metastasis. This study assessed the toxic effects of CuO NPs in endometrial and cervical cancer cell models as a potential anti-neoplastic agent for further integration into gynecological cancer drug delivery systems. Characterization of CuO NPs was used to confirm their size, shape, and charge. To explore the genotoxic effects, mechanisms of toxicity, and therapeutic potential of CuO NPs to target endometrial and cervical cancer, a palette of *in vitro* cellular assays were executed. To quantify the uptake of copper into cells, ICP-MS was performed on dry cell pellets revealing the relative amount of copper present in lysed cells. MTT assays exploring cell viability were used to generate IC_50_ curves and values, quantifying the sensitivity to CuO NPs for each cell line tested. To explore the CuO-NP’s potential to prevent the growth and spread of cancerous cells, cell migration assays were performed, revealing CuO NPs’ ability to inhibit cancer cell growth and spread, potentially mitigating the metastatic potential of these malignant cells. Oxidative stress analysis using ratios of GSH/GSSG revealed that all cell lines experienced significant changes to antioxidant potential after exposure to CuO NPs after only one hour of exposure, except for HeLa S3 *MLH1* KO.

## Future Perspectives

This work demonstrates critical discoveries in CuO NPs interactions with gynecological malignancies. Future work will focus on surface modification and interfacial engineering of these particles to enhance targeted delivery to cancer cells and improve colloidal stability in physiological environments. Alternatively, research also shows that combination therapies with antioxidants, such as curcumin, may be effective at mediating specific targeting of tumor mitochondria.^61^ In addition, this work will be validated in models with enhanced physiological relevance. To complement the 2D cell culture, emerging technologies with 3D mono- and co-culture,^62, 63^ as well as organ-on-a-chip,^64, 65^ provide robust, engineered non-animal models. Given the findings in this paper, that CuO NP efficacy varies across the model cell lines used, expanding the studies to a more comprehensive profile of oxidative stress, metabolic activity, and DNA damage represents a promising avenue for future discoveries.

## Acknowledgements

This publication was made possible by Grant Number K12DA035150 from the Office of Women’s Health Research and the National Institute on Drug Abuse at the National Institutes of Health (NIH) and Grant Number P30 ES026529 from the National Institute of Environmental Health Sciences at the National Institutes of Health. Its contents are solely the responsibility of the authors and do not necessarily represent the official views of NIH. This research was supported by the Oak Ridge Associated Universities Ralph E. Powe Junior Faculty Enhancement Award to B.E.G.

SEM, TEM, and EDX were performed at the University of Kentucky Electron Microscopy Center (EMC), a member of the National Nanotechnology Coordinated Infrastructure (NNCI), which is supported by the National Science Foundation (NNCI-2025075). Apoptosis was completed at the Flow Cytometry and Immune Monitoring Core (FCIM). ICP-MS was completed with the assistance of Dr. Jason M. Unrine and Shristi Shrestha with the University of Kentucky Center for Appalachian Research in Environmental Sciences (UK-CARES) in the Environmental Chemistry and Toxicology Laboratory.

Certain cell studies presented in this manuscript used the HeLa cell line. Henrietta Lacks, and the HeLa cell line that was established from her tumor cells without her knowledge or consent in 1951, have made significant contributions to scientific progress and advances in human health. We are grateful to Henrietta Lacks, now deceased, and to her surviving family members for their contributions to biomedical research.

## Author Contributions

Conceptualization, Project Administration, Supervision: B.G., E.G.

Data curation, Investigation, Methodology: J.B., C.R., L.M., B.K.

Funding Acquisition: B.G.

Validation, Visualization, Writing – original draft, Writing – review and editing: J.B., C.R., L.M. B. K., E.G., B.G.

