## Supporting Information for "Copper oxide nanoparticles function as antineoplastic agents in uterine cancer cell lines"

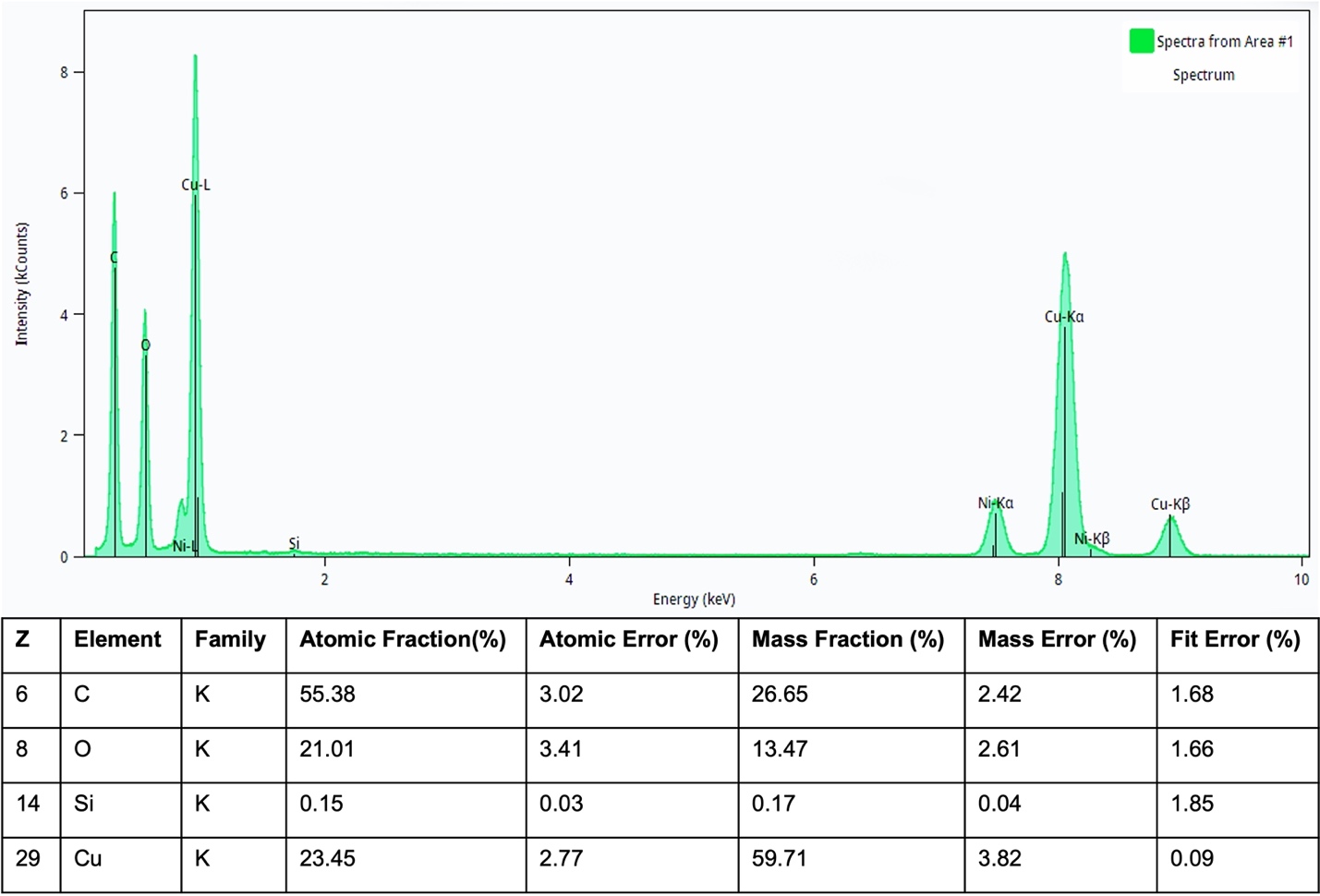

**Figure S1. EDS elemental composition of copper (Cu) and oxygen (O) in nanoparticles and carbon (C) and silicon (Si) from the background.**

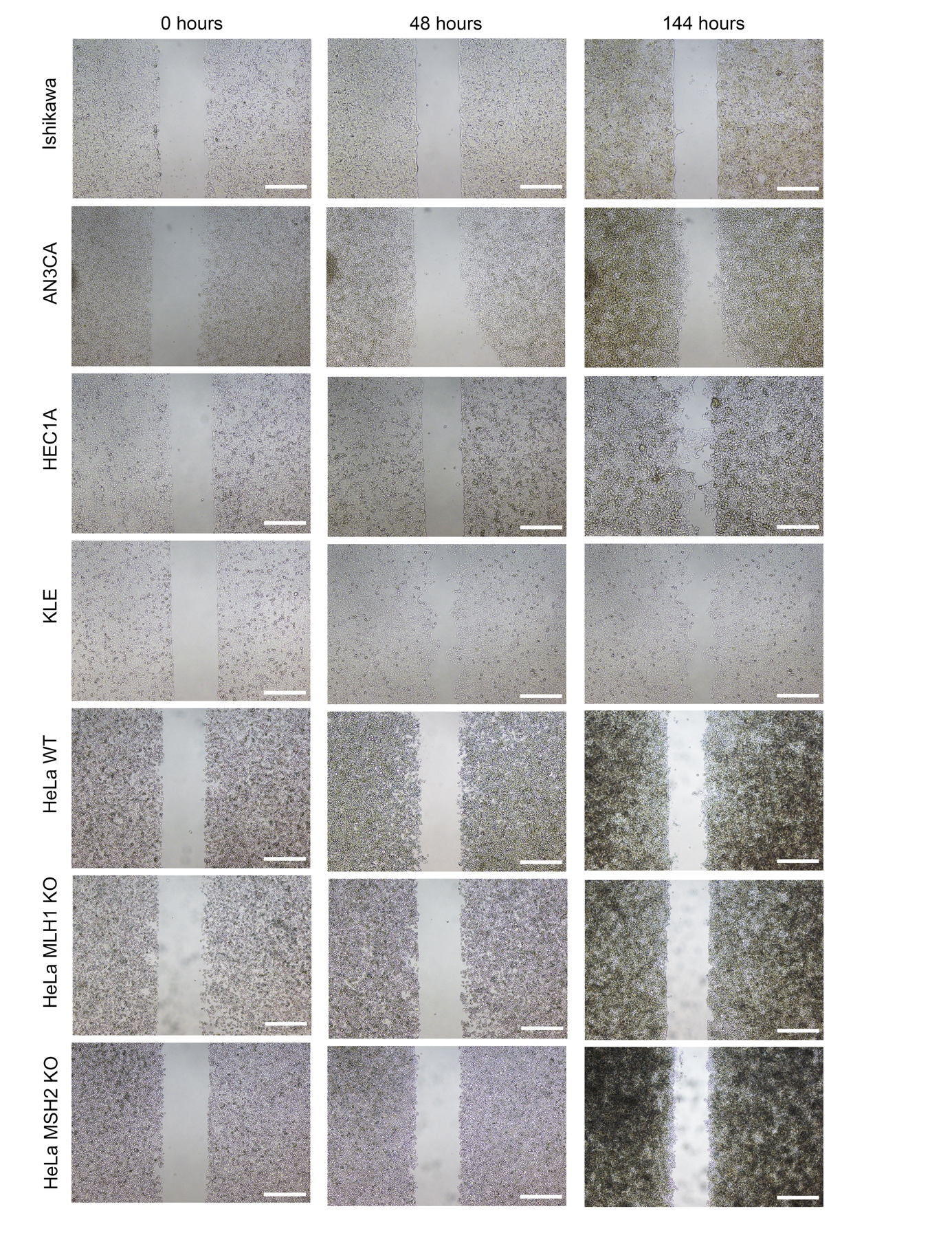
**Figure S2. Representative images of the scratch assay at 0 hours (t = 0 hours) for all cell lines with no CuO NP treatment.**

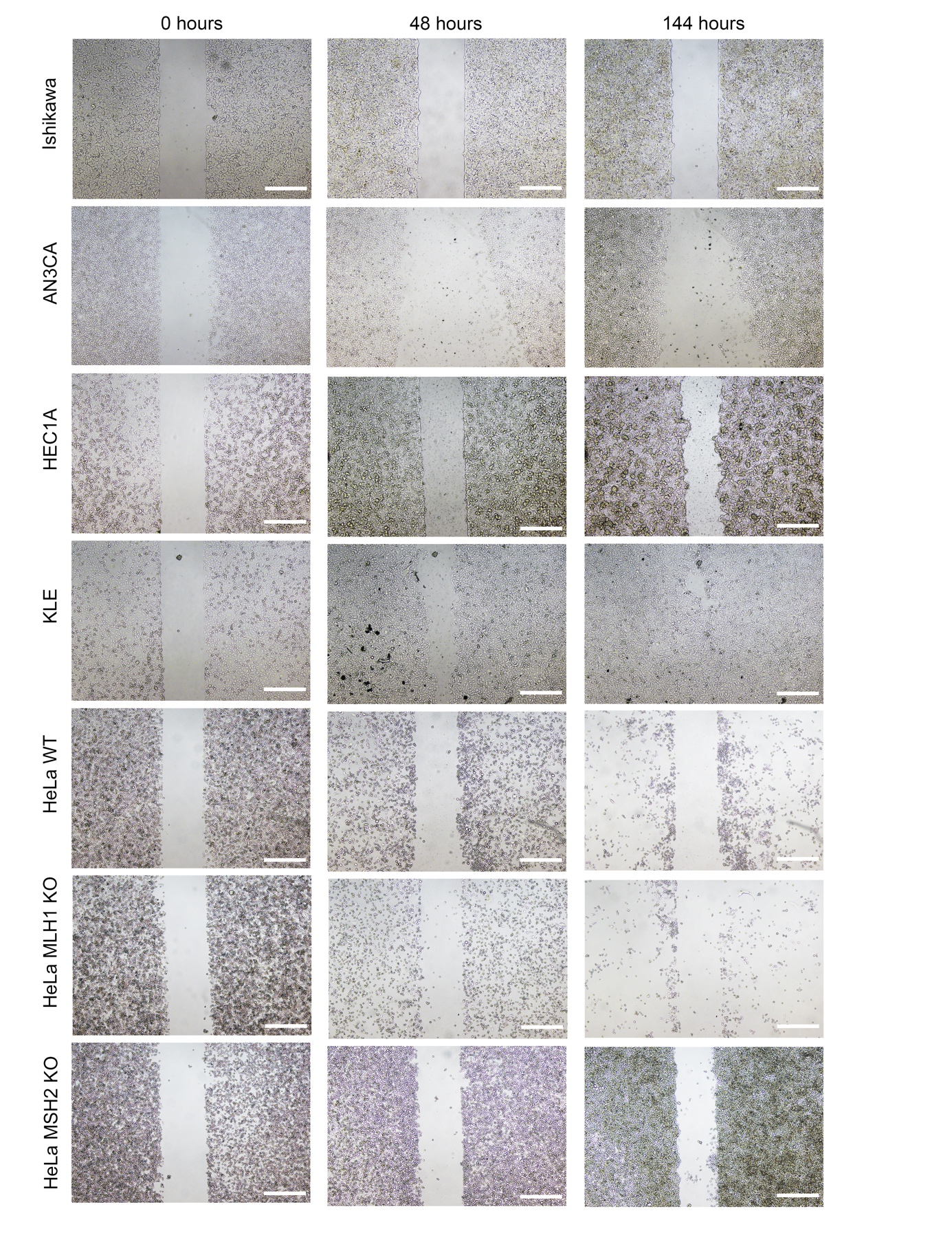

**Figure S3. Representative images of scratch assay at 48 hours (t = 48 hours) for all cell lines with 10 𝜇g/mL CuO NPs**

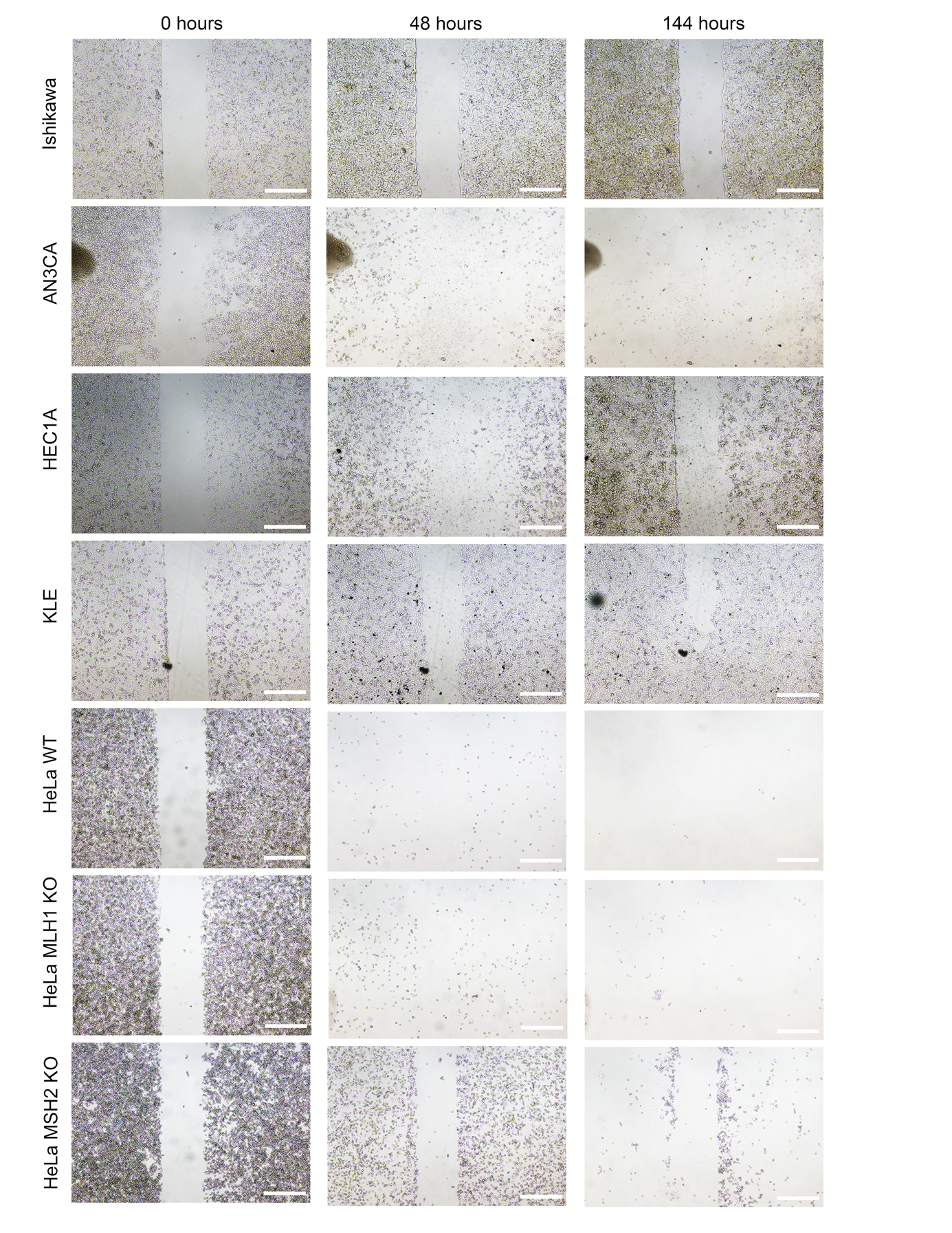
**Figure S4. Representative images of scratch assay at 144 hours (t = 144 hours) for all cell lines with 20 𝜇g/mL CuO NPs**

**Table S1. Statistical analysis of significant differences in intracellular copper (Cu^2+^) attained from ICP-MS. Comparisons were conducted using two-way ANOVA with Sidak’s multiple comparison.**

| Šídák's multiple comparisons test | Summary | Adjusted P Value |
| --- | --- | --- |
| 0 𝜇g/mL:AN3CA vs. 0 𝜇g/mL:Hec1A | ns | >0.9999 |
| 0 𝜇g/mL:AN3CA vs. 0 𝜇g/mL:KLE | ns | >0.9999 |
| 0 𝜇g/mL:AN3CA vs. 0 𝜇g/mL:HeLa S3 WT | ns | >0.9999 |
| 0 𝜇g/mL:AN3CA vs. 0 𝜇g/mL:HeLa S3 MLH1 KO | ns | >0.9999 |
| 0 𝜇g/mL:AN3CA vs. 0 𝜇g/mL:HeLa S3 MSH2 KO | ns | >0.9999 |
| 0 𝜇g/mL:AN3CA vs. 10 𝜇g/mL:AN3CA | ns | >0.9999 |
| 0 𝜇g/mL:AN3CA vs. 20 𝜇g/mL:AN3CA | ns | 0.492 |
| 0 𝜇g/mL:Hec1A vs. 0 𝜇g/mL:KLE | ns | >0.9999 |
| 0 𝜇g/mL:Hec1A vs. 0 𝜇g/mL:HeLa S3 WT | ns | >0.9999 |
| 0 𝜇g/mL:Hec1A vs. 0 𝜇g/mL:HeLa S3 MLH1 KO | ns | >0.9999 |
| 0 𝜇g/mL:Hec1A vs. 0 𝜇g/mL:HeLa S3 MSH2 KO | ns | >0.9999 |
| 0 𝜇g/mL:Hec1A vs. 10 𝜇g/mL:Ishikawa | *** | 0.0002 |
| 0 𝜇g/mL:Hec1A vs. 10 𝜇g/mL:AN3CA | ns | >0.9999 |
| 0 𝜇g/mL:Hec1A vs. 10 𝜇g/mL:Hec1A | ns | >0.9999 |
| 0 𝜇g/mL:Hec1A vs. 10 𝜇g/mL:KLE | ns | >0.9999 |
| 0 𝜇g/mL:Hec1A vs. 10 𝜇g/mL:HeLa S3 WT | ns | >0.9999 |
| 0 𝜇g/mL:Hec1A vs. 10 𝜇g/mL:HeLa S3 MLH1 KO | ns | >0.9999 |
| 0 𝜇g/mL:Hec1A vs. 10 𝜇g/mL:HeLa S3 MSH2 KO | ns | >0.9999 |
| 0 𝜇g/mL:Hec1A vs. 20 𝜇g/mL:Ishikawa | **** | <0.0001 |
| 0 𝜇g/mL:Hec1A vs. 20 𝜇g/mL:AN3CA | ns | >0.9999 |
| 0 𝜇g/mL:Hec1A vs. 20 𝜇g/mL:Hec1A | ns | 0.492 |
| 0 𝜇g/mL:Hec1A vs. 20 𝜇g/mL:KLE | ns | 0.9261 |
| 0 𝜇g/mL:Hec1A vs. 20 𝜇g/mL:HeLa S3 WT | ns | >0.9999 |
| 0 𝜇g/mL:Hec1A vs. 20 𝜇g/mL:HeLa S3 MLH1 KO | ns | >0.9999 |
| 0 𝜇g/mL:Hec1A vs. 20 𝜇g/mL:HeLa S3 MSH2 KO | ns | >0.9999 |
| 0 𝜇g/mL:KLE vs. 0 𝜇g/mL:HeLa S3 WT | ns | >0.9999 |
| 0 𝜇g/mL:KLE vs. 0 𝜇g/mL:HeLa S3 MLH1 KO | ns | >0.9999 |
| 0 𝜇g/mL:KLE vs. 0 𝜇g/mL:HeLa S3 MSH2 KO | ns | >0.9999 |
| 0 𝜇g/mL:KLE vs. 10 𝜇g/mL:Ishikawa | ** | 0.0026 |
| 0 𝜇g/mL:KLE vs. 10 𝜇g/mL:AN3CA | ns | >0.9999 |
| 0 𝜇g/mL:KLE vs. 10 𝜇g/mL:Hec1A | ns | >0.9999 |
| 0 𝜇g/mL:KLE vs. 10 𝜇g/mL:KLE | ns | >0.9999 |
| 0 𝜇g/mL:KLE vs. 10 𝜇g/mL:HeLa S3 WT | ns | >0.9999 |
| 0 𝜇g/mL:KLE vs. 10 𝜇g/mL:HeLa S3 MLH1 KO | ns | >0.9999 |
| 0 𝜇g/mL:KLE vs. 10 𝜇g/mL:HeLa S3 MSH2 KO | ns | >0.9999 |
| 0 𝜇g/mL:KLE vs. 20 𝜇g/mL:Ishikawa | **** | <0.0001 |
| 0 𝜇g/mL:KLE vs. 20 𝜇g/mL:AN3CA | ns | >0.9999 |
| 0 𝜇g/mL:KLE vs. 20 𝜇g/mL:Hec1A | ns | >0.9999 |
| 0 𝜇g/mL:KLE vs. 20 𝜇g/mL:KLE | ns | 0.492 |
| 0 𝜇g/mL:KLE vs. 20 𝜇g/mL:HeLa S3 WT | ns | >0.9999 |
| 0 𝜇g/mL:KLE vs. 20 𝜇g/mL:HeLa S3 MLH1 KO | ns | >0.9999 |
| 0 𝜇g/mL:KLE vs. 20 𝜇g/mL:HeLa S3 MSH2 KO | ns | >0.9999 |
| 0 𝜇g/mL:HeLa S3 WT vs. 0 𝜇g/mL:HeLa S3 MLH1 KO | ns | >0.9999 |
| 0 𝜇g/mL:HeLa S3 WT vs. 0 𝜇g/mL:HeLa S3 MSH2 KO | ns | >0.9999 |
| 0 𝜇g/mL:HeLa S3 WT vs. 10 𝜇g/mL:Ishikawa | *** | 0.0002 |
| 0 𝜇g/mL:HeLa S3 WT vs. 10 𝜇g/mL:AN3CA | ns | >0.9999 |
| 0 𝜇g/mL:HeLa S3 WT vs. 10 𝜇g/mL:Hec1A | ns | >0.9999 |
| 0 𝜇g/mL:HeLa S3 WT vs. 10 𝜇g/mL:KLE | ns | >0.9999 |
| 0 𝜇g/mL:HeLa S3 WT vs. 10 𝜇g/mL:HeLa S3 WT | ns | >0.9999 |
| 0 𝜇g/mL:HeLa S3 WT vs. 10 𝜇g/mL:HeLa S3 MLH1 KO | ns | >0.9999 |
| 0 𝜇g/mL:HeLa S3 WT vs. 10 𝜇g/mL:HeLa S3 MSH2 KO | ns | >0.9999 |
| 0 𝜇g/mL:HeLa S3 WT vs. 20 𝜇g/mL:Ishikawa | **** | <0.0001 |
| 0 𝜇g/mL:HeLa S3 WT vs. 20 𝜇g/mL:AN3CA | ns | >0.9999 |
| 0 𝜇g/mL:HeLa S3 WT vs. 20 𝜇g/mL:Hec1A | ns | >0.9999 |
| 0 𝜇g/mL:HeLa S3 WT vs. 20 𝜇g/mL:KLE | ns | 0.9083 |
| 0 𝜇g/mL:HeLa S3 WT vs. 20 𝜇g/mL:HeLa S3 WT | ns | 0.492 |
| 0 𝜇g/mL:HeLa S3 WT vs. 20 𝜇g/mL:HeLa S3 MLH1 KO | ns | >0.9999 |
| 0 𝜇g/mL:HeLa S3 WT vs. 20 𝜇g/mL:HeLa S3 MSH2 KO | ns | >0.9999 |
| 0 𝜇g/mL:HeLa S3 MLH1 KO vs. 0 𝜇g/mL:HeLa S3 MSH2 KO | ns | >0.9999 |
| 0 𝜇g/mL:HeLa S3 MLH1 KO vs. 10 𝜇g/mL:Ishikawa | **** | <0.0001 |
| 0 𝜇g/mL:HeLa S3 MLH1 KO vs. 10 𝜇g/mL:AN3CA | ns | >0.9999 |
| 0 𝜇g/mL:HeLa S3 MLH1 KO vs. 10 𝜇g/mL:Hec1A | ns | >0.9999 |
| 0 𝜇g/mL:HeLa S3 MLH1 KO vs. 10 𝜇g/mL:KLE | ns | >0.9999 |
| 0 𝜇g/mL:HeLa S3 MLH1 KO vs. 10 𝜇g/mL:HeLa S3 WT | ns | >0.9999 |
| 0 𝜇g/mL:HeLa S3 MLH1 KO vs. 10 𝜇g/mL:HeLa S3 MLH1 KO | ns | >0.9999 |
| 0 𝜇g/mL:HeLa S3 MLH1 KO vs. 10 𝜇g/mL:HeLa S3 MSH2 KO | ns | >0.9999 |
| 0 𝜇g/mL:HeLa S3 MLH1 KO vs. 20 𝜇g/mL:Ishikawa | **** | <0.0001 |
| 0 𝜇g/mL:HeLa S3 MLH1 KO vs. 20 𝜇g/mL:AN3CA | ns | >0.9999 |
| 0 𝜇g/mL:HeLa S3 MLH1 KO vs. 20 𝜇g/mL:Hec1A | ns | >0.9999 |
| 0 𝜇g/mL:HeLa S3 MLH1 KO vs. 20 𝜇g/mL:KLE | ns | 0.7253 |
| 0 𝜇g/mL:HeLa S3 MLH1 KO vs. 20 𝜇g/mL:HeLa S3 WT | ns | >0.9999 |
| 0 𝜇g/mL:HeLa S3 MLH1 KO vs. 20 𝜇g/mL:HeLa S3 MLH1 KO | ns | 0.492 |
| 0 𝜇g/mL:HeLa S3 MLH1 KO vs. 20 𝜇g/mL:HeLa S3 MSH2 KO | ns | >0.9999 |
| 0 𝜇g/mL:HeLa S3 MSH2 KO vs. 10 𝜇g/mL:Ishikawa | **** | <0.0001 |
| 0 𝜇g/mL:HeLa S3 MSH2 KO vs. 10 𝜇g/mL:AN3CA | ns | >0.9999 |
| 0 𝜇g/mL:HeLa S3 MSH2 KO vs. 10 𝜇g/mL:Hec1A | ns | >0.9999 |
| 0 𝜇g/mL:HeLa S3 MSH2 KO vs. 10 𝜇g/mL:KLE | ns | >0.9999 |
| 0 𝜇g/mL:HeLa S3 MSH2 KO vs. 10 𝜇g/mL:HeLa S3 WT | ns | >0.9999 |
| 0 𝜇g/mL:HeLa S3 MSH2 KO vs. 10 𝜇g/mL:HeLa S3 MLH1 KO | ns | >0.9999 |
| 0 𝜇g/mL:HeLa S3 MSH2 KO vs. 10 𝜇g/mL:HeLa S3 MSH2 KO | ns | >0.9999 |
| 0 𝜇g/mL:HeLa S3 MSH2 KO vs. 20 𝜇g/mL:Ishikawa | **** | <0.0001 |
| 0 𝜇g/mL:HeLa S3 MSH2 KO vs. 20 𝜇g/mL:AN3CA | ns | 0.9992 |
| 0 𝜇g/mL:HeLa S3 MSH2 KO vs. 20 𝜇g/mL:Hec1A | ns | 0.9987 |
| 0 𝜇g/mL:HeLa S3 MSH2 KO vs. 20 𝜇g/mL:KLE | ns | 0.5838 |
| 0 𝜇g/mL:HeLa S3 MSH2 KO vs. 20 𝜇g/mL:HeLa S3 WT | ns | 0.9993 |
| 0 𝜇g/mL:HeLa S3 MSH2 KO vs. 20 𝜇g/mL:HeLa S3 MLH1 KO | ns | >0.9999 |
| 0 𝜇g/mL:HeLa S3 MSH2 KO vs. 20 𝜇g/mL:HeLa S3 MSH2 KO | ns | 0.492 |
| 10 𝜇g/mL:Ishikawa  vs. 10 𝜇g/mL:AN3CA | **** | <0.0001 |
| 10 𝜇g/mL:Ishikawa  vs. 10 𝜇g/mL:Hec1A | **** | <0.0001 |
| 10 𝜇g/mL:Ishikawa  vs. 10 𝜇g/mL:KLE | ** | 0.0011 |
| 10 𝜇g/mL:Ishikawa  vs. 10 𝜇g/mL:HeLa S3 WT | **** | <0.0001 |
| 10 𝜇g/mL:Ishikawa  vs. 10 𝜇g/mL:HeLa S3 MLH1 KO | **** | <0.0001 |
| 10 𝜇g/mL:Ishikawa  vs. 10 𝜇g/mL:HeLa S3 MSH2 KO | **** | <0.0001 |
| 10 𝜇g/mL:Ishikawa  vs. 20 𝜇g/mL:Ishikawa | ns | 0.5615 |
| 10 𝜇g/mL:Ishikawa  vs. 20 𝜇g/mL:AN3CA | ns | 0.1221 |
| 10 𝜇g/mL:Ishikawa  vs. 20 𝜇g/mL:Hec1A | ns | 0.1162 |
| 10 𝜇g/mL:Ishikawa  vs. 20 𝜇g/mL:KLE | ns | 0.6529 |
| 10 𝜇g/mL:Ishikawa  vs. 20 𝜇g/mL:HeLa S3 WT | ns | 0.106 |
| 10 𝜇g/mL:Ishikawa  vs. 20 𝜇g/mL:HeLa S3 MLH1 KO | ns | 0.0552 |
| 10 𝜇g/mL:Ishikawa  vs. 20 𝜇g/mL:HeLa S3 MSH2 KO | * | 0.0366 |
| 10 𝜇g/mL:AN3CA vs. 10 𝜇g/mL:Hec1A | ns | >0.9999 |
| 10 𝜇g/mL:AN3CA vs. 10 𝜇g/mL:KLE | ns | >0.9999 |
| 10 𝜇g/mL:AN3CA vs. 10 𝜇g/mL:HeLa S3 WT | ns | >0.9999 |
| 10 𝜇g/mL:AN3CA vs. 10 𝜇g/mL:HeLa S3 MLH1 KO | ns | >0.9999 |
| 10 𝜇g/mL:AN3CA vs. 10 𝜇g/mL:HeLa S3 MSH2 KO | ns | >0.9999 |
| 10 𝜇g/mL:AN3CA vs. 20 𝜇g/mL:Ishikawa | **** | <0.0001 |
| 10 𝜇g/mL:AN3CA vs. 20 𝜇g/mL:AN3CA | ns | 0.5615 |
| 10 𝜇g/mL:AN3CA vs. 20 𝜇g/mL:Hec1A | ns | >0.9999 |
| 10 𝜇g/mL:AN3CA vs. 20 𝜇g/mL:KLE | ns | 0.9736 |
| 10 𝜇g/mL:AN3CA vs. 20 𝜇g/mL:HeLa S3 WT | ns | >0.9999 |
| 10 𝜇g/mL:AN3CA vs. 20 𝜇g/mL:HeLa S3 MLH1 KO | ns | >0.9999 |
| 10 𝜇g/mL:AN3CA vs. 20 𝜇g/mL:HeLa S3 MSH2 KO | ns | >0.9999 |
| 10 𝜇g/mL:Hec1A vs. 10 𝜇g/mL:KLE | ns | >0.9999 |
| 10 𝜇g/mL:Hec1A vs. 10 𝜇g/mL:HeLa S3 WT | ns | >0.9999 |
| 10 𝜇g/mL:Hec1A vs. 10 𝜇g/mL:HeLa S3 MLH1 KO | ns | >0.9999 |
| 10 𝜇g/mL:Hec1A vs. 10 𝜇g/mL:HeLa S3 MSH2 KO | ns | >0.9999 |
| 10 𝜇g/mL:Hec1A vs. 20 𝜇g/mL:Ishikawa | **** | <0.0001 |
| 10 𝜇g/mL:Hec1A vs. 20 𝜇g/mL:AN3CA | ns | >0.9999 |
| 10 𝜇g/mL:Hec1A vs. 20 𝜇g/mL:Hec1A | ns | 0.5615 |
| 10 𝜇g/mL:Hec1A vs. 20 𝜇g/mL:KLE | ns | 0.9674 |
| 10 𝜇g/mL:Hec1A vs. 20 𝜇g/mL:HeLa S3 WT | ns | >0.9999 |
| 10 𝜇g/mL:Hec1A vs. 20 𝜇g/mL:HeLa S3 MLH1 KO | ns | >0.9999 |
| 10 𝜇g/mL:Hec1A vs. 20 𝜇g/mL:HeLa S3 MSH2 KO | ns | >0.9999 |
| 10 𝜇g/mL:KLE vs. 10 𝜇g/mL:HeLa S3 WT | ns | >0.9999 |
| 10 𝜇g/mL:KLE vs. 10 𝜇g/mL:HeLa S3 MLH1 KO | ns | >0.9999 |
| 10 𝜇g/mL:KLE vs. 10 𝜇g/mL:HeLa S3 MSH2 KO | ns | >0.9999 |
| 10 𝜇g/mL:KLE vs. 20 𝜇g/mL:Ishikawa | **** | <0.0001 |
| 10 𝜇g/mL:KLE vs. 20 𝜇g/mL:AN3CA | ns | >0.9999 |
| 10 𝜇g/mL:KLE vs. 20 𝜇g/mL:Hec1A | ns | >0.9999 |
| 10 𝜇g/mL:KLE vs. 20 𝜇g/mL:KLE | ns | 0.5615 |
| 10 𝜇g/mL:KLE vs. 20 𝜇g/mL:HeLa S3 WT | ns | >0.9999 |
| 10 𝜇g/mL:KLE vs. 20 𝜇g/mL:HeLa S3 MLH1 KO | ns | >0.9999 |
| 10 𝜇g/mL:KLE vs. 20 𝜇g/mL:HeLa S3 MSH2 KO | ns | >0.9999 |
| 10 𝜇g/mL:HeLa S3 WT vs. 10 𝜇g/mL:HeLa S3 MLH1 KO | ns | >0.9999 |
| 10 𝜇g/mL:HeLa S3 WT vs. 10 𝜇g/mL:HeLa S3 MSH2 KO | ns | >0.9999 |
| 10 𝜇g/mL:HeLa S3 WT vs. 20 𝜇g/mL:Ishikawa | **** | <0.0001 |
| 10 𝜇g/mL:HeLa S3 WT vs. 20 𝜇g/mL:AN3CA | ns | >0.9999 |
| 10 𝜇g/mL:HeLa S3 WT vs. 20 𝜇g/mL:Hec1A | ns | >0.9999 |
| 10 𝜇g/mL:HeLa S3 WT vs. 20 𝜇g/mL:KLE | ns | 0.9567 |
| 10 𝜇g/mL:HeLa S3 WT vs. 20 𝜇g/mL:HeLa S3 WT | ns | 0.5615 |
| 10 𝜇g/mL:HeLa S3 WT vs. 20 𝜇g/mL:HeLa S3 MLH1 KO | ns | >0.9999 |
| 10 𝜇g/mL:HeLa S3 WT vs. 20 𝜇g/mL:HeLa S3 MSH2 KO | ns | >0.9999 |
| 10 𝜇g/mL:HeLa S3 MLH1 KO vs. 10 𝜇g/mL:HeLa S3 MSH2 KO | ns | >0.9999 |
| 10 𝜇g/mL:HeLa S3 MLH1 KO vs. 20 𝜇g/mL:Ishikawa | **** | <0.0001 |
| 10 𝜇g/mL:HeLa S3 MLH1 KO vs. 20 𝜇g/mL:AN3CA | ns | >0.9999 |
| 10 𝜇g/mL:HeLa S3 MLH1 KO vs. 20 𝜇g/mL:Hec1A | ns | >0.9999 |
| 10 𝜇g/mL:HeLa S3 MLH1 KO vs. 20 𝜇g/mL:KLE | ns | 0.8164 |
| 10 𝜇g/mL:HeLa S3 MLH1 KO vs. 20 𝜇g/mL:HeLa S3 WT | ns | >0.9999 |
| 10 𝜇g/mL:HeLa S3 MLH1 KO vs. 20 𝜇g/mL:HeLa S3 MLH1 KO | ns | 0.5615 |
| 10 𝜇g/mL:HeLa S3 MLH1 KO vs. 20 𝜇g/mL:HeLa S3 MSH2 KO | ns | >0.9999 |
| 10 𝜇g/mL:HeLa S3 MSH2 KO vs. 20 𝜇g/mL:Ishikawa | **** | <0.0001 |
| 10 𝜇g/mL:HeLa S3 MSH2 KO vs. 20 𝜇g/mL:AN3CA | ns | >0.9999 |
| 10 𝜇g/mL:HeLa S3 MSH2 KO vs. 20 𝜇g/mL:Hec1A | ns | 0.9998 |
| 10 𝜇g/mL:HeLa S3 MSH2 KO vs. 20 𝜇g/mL:KLE | ns | 0.6826 |
| 10 𝜇g/mL:HeLa S3 MSH2 KO vs. 20 𝜇g/mL:HeLa S3 WT | ns | >0.9999 |
| 10 𝜇g/mL:HeLa S3 MSH2 KO vs. 20 𝜇g/mL:HeLa S3 MLH1 KO | ns | >0.9999 |
| 10 𝜇g/mL:HeLa S3 MSH2 KO vs. 20 𝜇g/mL:HeLa S3 MSH2 KO | ns | 0.5615 |
| 20 𝜇g/mL:Ishikawa  vs. 20 𝜇g/mL:AN3CA | **** | <0.0001 |
| 20 𝜇g/mL:Ishikawa  vs. 20 𝜇g/mL:Hec1A | **** | <0.0001 |
| 20 𝜇g/mL:Ishikawa  vs. 20 𝜇g/mL:KLE | ** | 0.0011 |
| 20 𝜇g/mL:Ishikawa  vs. 20 𝜇g/mL:HeLa S3 WT | **** | <0.0001 |
| 20 𝜇g/mL:Ishikawa  vs. 20 𝜇g/mL:HeLa S3 MLH1 KO | **** | <0.0001 |
| 20 𝜇g/mL:Ishikawa  vs. 20 𝜇g/mL:HeLa S3 MSH2 KO | **** | <0.0001 |
| 20 𝜇g/mL:AN3CA vs. 20 𝜇g/mL:Hec1A | ns | >0.9999 |
| 20 𝜇g/mL:AN3CA vs. 20 𝜇g/mL:KLE | ns | >0.9999 |
| 20 𝜇g/mL:AN3CA vs. 20 𝜇g/mL:HeLa S3 WT | ns | >0.9999 |
| 20 𝜇g/mL:AN3CA vs. 20 𝜇g/mL:HeLa S3 MLH1 KO | ns | >0.9999 |
| 20 𝜇g/mL:AN3CA vs. 20 𝜇g/mL:HeLa S3 MSH2 KO | ns | >0.9999 |
| 20 𝜇g/mL:Hec1A vs. 20 𝜇g/mL:KLE | ns | >0.9999 |
| 20 𝜇g/mL:Hec1A vs. 20 𝜇g/mL:HeLa S3 WT | ns | >0.9999 |
| 20 𝜇g/mL:Hec1A vs. 20 𝜇g/mL:HeLa S3 MLH1 KO | ns | >0.9999 |
| 20 𝜇g/mL:Hec1A vs. 20 𝜇g/mL:HeLa S3 MSH2 KO | ns | >0.9999 |
| 20 𝜇g/mL:KLE vs. 20 𝜇g/mL:HeLa S3 WT | ns | >0.9999 |
| 20 𝜇g/mL:KLE vs. 20 𝜇g/mL:HeLa S3 MLH1 KO | ns | >0.9999 |
| 20 𝜇g/mL:KLE vs. 20 𝜇g/mL:HeLa S3 MSH2 KO | ns | >0.9999 |
| 20 𝜇g/mL:HeLa S3 WT vs. 20 𝜇g/mL:HeLa S3 MLH1 KO | ns | >0.9999 |
| 20 𝜇g/mL:HeLa S3 WT vs. 20 𝜇g/mL:HeLa S3 MSH2 KO | ns | >0.9999 |
| 20 𝜇g/mL:HeLa S3 MLH1 KO vs. 20 𝜇g/mL:HeLa S3 MSH2 KO | ns | >0.9999 |
